# Restriction of innate Tγδ17 cell plasticity by an AP-1 regulatory axis

**DOI:** 10.1101/2024.10.15.618522

**Authors:** Morgan E. Parker, Naren U. Mehta, Tzu-Chieh Liao, William H. Tomaszewski, Stephanie A. Snyder, Julia Busch, Maria Ciofani

**Affiliations:** Department of Integrative Immunobiology, Duke University Medical Center, Durham, NC, USA; Center for Advanced Genomic Technologies, Duke University, Durham, NC, USA; Department of Molecular Genetics and Microbiology, Duke University Medical Center, Durham, NC, USA

## Abstract

IL-17-producing γδ T (Tγδ17) cells are innate-like mediators of intestinal barrier immunity. While Th17 cell and ILC3 plasticity have been extensively studied, the mechanisms governing Tγδ17 cell effector flexibility remain undefined. Here, we combined type 3 fate-mapping with single cell ATAC/RNA-seq multiome profiling to define the cellular features and regulatory networks underlying Tγδ17 cell plasticity. During homeostasis, Tγδ17 cell effector identity was stable across tissues, including for intestinal T-bet^+^ Tγδ17 cells that restrained IFNγ production. However, S. typhimurium infection induced intestinal Vγ6^+^ Tγδ17 cell conversion into type 1 effectors, with loss of IL-17A production and partial RORγt downregulation. Multiome analysis revealed a trajectory along Vγ6^+^ Tγδ17 effector conversion, with TIM-3 marking ex-Tγδ17 cells with enhanced type 1 functionality. Lastly, we characterized and validated a critical AP-1 regulatory axis centered around JunB and Fosl2 that controls Vγ6^+^ Tγδ17 cell plasticity by stabilizing type 3 identity and restricting type 1 effector conversion.

## Introduction

Innate-like γδ T cells are unconventional T cells that share features with both innate and adaptive lymphocytes^1–4^. Although innate-like γδ T cells express T cell receptors (TCRs), albeit with limited diversity, their effector fate is programmed in the thymus during ontogeny, facilitating immediate responses after cytokine stimulation^5,6^. As tissue-resident lymphocytes, γδ T cells sense their local environment and are regulated through a combination of the TCR, cytokine receptors, co-stimulatory receptors, inhibitory receptors, and natural killer receptors^7,8^. Given their widespread distribution, enrichment at barrier tissues, and early seeding during fetal life, innate-like γδ T cells act as a first line of defense for tissue-sensing and protection against invading pathogens^1^. Thus, γδ T cells play numerous roles in tissue homeostasis, wound healing, and pathogen and tumor clearance^1^. Although protective in many settings, unrestrained γδ T cells contribute to chronic inflammation and autoimmune diseases^9^.

Broadly, γδ T cells can be divided into two main effector subsets based on cytokine and lineage defining transcription factor (LDTF) expression: type 3 IL-17A-producers expressing RORγt (Tγδ17 cells) and type 1 IFNγ-producers expressing T-bet (Tγδ1 cells)^6^. IL-17A production by γδ T cells is linked to clearance of extracellular bacteria and fungi, while IFNγ production by γδ T cells is associated with anti-tumor responses and clearance of intracellular pathogens^5^. The Tγδ17 cell compartment is comprised of cells expressing TCRs with Vγ6 or Vγ4 γ chain variable regions, with each subset having distinct ontogenies (Tonegawa nomenclature)^10,11^. Vγ6^+^ Tγδ17 cell development is restricted to a fetal wave in the thymus between E16 and E18 and cannot be reconstituted with adult bone marrow^11^. In contrast, Vγ4^+^ Tγδ17 cells are heterogenous in that one subset develops from a perinatal wave as innate-like Tγδ17 cells, while others exit the thymus as naïve cells and are ‘inducible’ Tγδ17 cells following activation and type 3 polarization in the periphery, akin to conventional αβ T cells^11,12^.

Innate and adaptive type 3 lymphocytes expressing RORγt—including ILC3s, Tγδ17 cells, and Th17 cells—are epigenetically primed for *Tbx21* (T-bet) and *Ifng* expression, leading to type 1 functional plasticity in certain settings^13–17^. For example, Tγδ17 cells in homeostatic conditions have permissive H3K4me2 modifications at the *Ifng* locus, while Tγδ1 cells display extensive repressive H3K27me3 marks at type 3 loci (*Rorc*, *Il17a*, *Blk*, *Maf*), suggesting that Tγδ17 cells are poised for alternative effector fates relative to their Tγδ1 counterparts^15^. Environmental requirements for type 1 plasticity are cell-type dependent, as ILC3s convert to ILC1-like cells at steady state^18^, while adaptive Th17 and IL-17A-producing CD8^+^ T (Tc17) cells require an inflammatory setting to convert to type 1 effectors^19–22^. Although such plasticity provides flexibility during infections, dysregulation of this process drives inflammation that is implicated in many autoimmune diseases, such as inflammatory bowel disease and multiple sclerosis^22–27^.

In contrast to the effector flexibility of type 3 ILCs and αβ T cells, our understanding of Tγδ17 cell plasticity remains limited. After short-term IL-23 and IL-1β stimulation, Tγδ17 cells robustly produce IL-17A, however, prolonged culture with these cytokines results in polyfunctional Tγδ17 cells that co-produce IL-17A and IFNγ^15,28^. Furthermore, such functional plasticity of Tγδ17 cells appears to be context-dependent, as IL-17A^+^IFNγ^+^ cells are only observed in specific inflammatory settings, such as oral *Listeria monocytogenes* infection^29^ and an ovarian cancer model^15^, but not in naïve mice nor during any systemic infections tested^15^. While these seminal studies have demonstrated the ability of Tγδ17 cells to co-produce IL-17A and IFNγ under conditions of strong inflammation, type 3 fate-mapping approaches were not employed to determine whether Tγδ17 cells have the capacity for full effector conversion to Tγδ1-like cells via loss of type 3 features.

The transcriptional regulators dictating the effector fate of developing γδ T cells have been thoroughly studied^5^, however, the key regulators of Tγδ17 cell plasticity remain largely unknown outside of a few studies. In particular, retinoic acid receptor α (RARα) signaling promotes the production of IFNγ by Tγδ17 cells in the intestine^30^, while the microRNA miR-146a post-transcriptionally restricts the generation of IL-17A^+^IFNγ^+^ Tγδ17 cells both *in vitro* and after *L. monocytogenes* infection via targeting of *Nod1*^31^. Regulating the flexibility of Tγδ17 cells may be clinically useful. Indeed, Tγδ17 and Tγδ1 cells have opposing roles in the context of cancer, with Tγδ17 cells promoting colorectal cancer progression through IL-17A production, while Tγδ1 cells exhibit anti-tumor properties^32^. Therefore, identifying the molecular switches governing type 1 plasticity in Tγδ17 cells could be advantageous in designing therapies for disease contexts that benefit from diverting cells away from IL-17A production and towards IFNγ production.

To further our understanding of Tγδ17 cell plasticity, we used type 3 fate-mapping in combination with single cell ATAC/RNA-seq multiome epigenomic and gene expression profiling to track and characterize Tγδ17 cell plasticity at the cellular and molecular level. Here, we report that in a variety of lymphoid and nonlymphoid tissues, the effector identity of fate-mapped Tγδ17 cells was stable at steady state. This includes intestinal Tγδ17 cells that co-express RORγt and T-bet^30^ that were functionally restrained with respect to IFNγ production. Notably, oral *Salmonella typhimurium* infection induced type 1 effector conversion of intestinal Tγδ17 cells into IFNγ single producers selectively in the innate-like Vγ6^+^ subset. Vγ6^+^ Tγδ17 cell plasticity was characterized by a loss of IL-17A production and an intermediate downregulation of RORγt expression, along with a global shift from a type 3 to type 1 effector expression program. Additionally, expression of the coinhibitory receptor TIM-3 selectively marked converted ex-Tγδ17 cells with enhanced type 1 functionality. An integrated analysis of gene expression, chromatin accessibility, TF motif enrichment, and gene regulatory networks that were altered along the single cell Vγ6^+^ Tγδ17 type 1 conversion trajectory revealed the dominant activity of a bZIP/AP-1 regulatory axis in this process. Among bZIP factors, we validated novel roles for AP-1 TFs JunB and Fosl2 in restricting intestinal Tγδ17 cell plasticity by stabilizing the expression of type 3 effectors (RORγt and IL-17A) and repressing the expression of type 1 effectors (T-bet or IFNγ). Genetic downmodulation of RORγt expression was sufficient for de-repression of IFNγ expression in Tγδ17 cells, thus identifying the regulation of RORγt as a critical node in the AP-1 network governing Vγ6^+^ Tγδ17 cell effector plasticity.

## Results

### Tγδ17 cell identity is stable at steady state

To investigate Tγδ17 cell plasticity, we employed two type 3 lymphocyte fate-mapping approaches that make use of mice harboring a conditional loxp-STOP-loxp-*ZsGreen* allele (*ZSG*) knocked into the *Rosa26* locus (*R26^ZSG^*)^33^ and bred to the *Rorc*-Cre or *Il17a*^Cre^ deleter strains^22,34^. The resulting *Rorc*-Cre *R26^ZSG^* and *Il17a*^Cre^ *R26^ZSG^* models mark all γδ T cells that have a history of *Rorc* and *Il17a* expression, respectively, with ZsGreen (ZS) expression. Thus, if Tγδ17 cells undergo type 1 lineage conversion into Tγδ1-like cells, such ‘ex-Tγδ17’ cells can be delineated from *bona fide* Tγδ1 cells. Importantly, both *Rorc*-Cre *R26^ZSG^*and *Il17a*^Cre^ *R26^ZSG^* models faithfully labeled Tγδ17 cells; in the colon, all RORγt^+^ γδ T cells expressed ZS due to prior *Rorc* or *Il17a* expression in the thymus during ontogeny (Extended Data Fig. 1a). Unlike the incomplete *Il17a* fate-mapping of Th17 cells^22^, there were no ZS^−^ IL-17A^+^ cells within the γδ T cell compartment of either model (Extended Data Fig. 1a). In the colon, Tγδ17 cells comprise Vγ4^+^ and Vγ6^+^ subsets (Extended Data Fig. 1b). Vγ6^+^ γδ T cells are uniformly ZS^+^ Tγδ17 cells, whereas Vγ4^+^ γδ T cells are a heterogenous population composed of both ZS^−^ Tγδ1 and ZS^+^ Tγδ17 cells (Extended Data Fig. 1b). Therefore, fate-mapping Tγδ17 cells facilitates the interrogation of Vγ4^+^ versus Vγ6^+^ Tγδ17 cell plasticity.

First, we assessed Tγδ17 cell lineage stability in *Rorc*-Cre *R26^ZSG^* mice. Of note, CD27 and CCR6 expression can be used in lymphoid tissues to distinguish Tγδ1 and Tγδ17 cells^35,36^, respectively, however, certain tissue environments and inflammation result in CCR6 downregulation^37^, and enzymes commonly used in tissue digestion cleave CD27^38^. Therefore, we relied on LDTF RORγt staining combined with fate-mapping to define γδ T cell populations; ZS^+^ RORγt^+^ γδ T cells represent current Tγδ17 cells and in the event of lineage conversion, ZS^+^ RORγt^−^ γδ T cells would mark ex-Tγδ17 cells. In both lymphoid (inguinal LN (iLN) and mLN) and nonlymphoid (small intestine lamina propria (siLP), colonic lamina propria (coLP), lung, and female reproductive tract (FRT)) tissues assayed, fate-mapped ZS^+^ γδ T cells uniformly expressed high levels of RORγt (Fig. 1a), indicating Tγδ17 cells maintain lineage identity during homeostasis. Next, we determined whether fate-mapped Tγδ17 cells exhibited functional plasticity in naive mice. Similar to the stability of RORγt expression, ZS^+^ γδ T cells in naive mice were robust IL-17A producers after ex vivo PMA/ionomycin stimulation and did not produce IFNγ, irrespective of the tissue assayed (Fig. 1b, c). These findings indicate that Tγδ17 cell identity is stable at steady state, both in terms of LDTF expression and functional cytokine production.

**Fig. 1.**
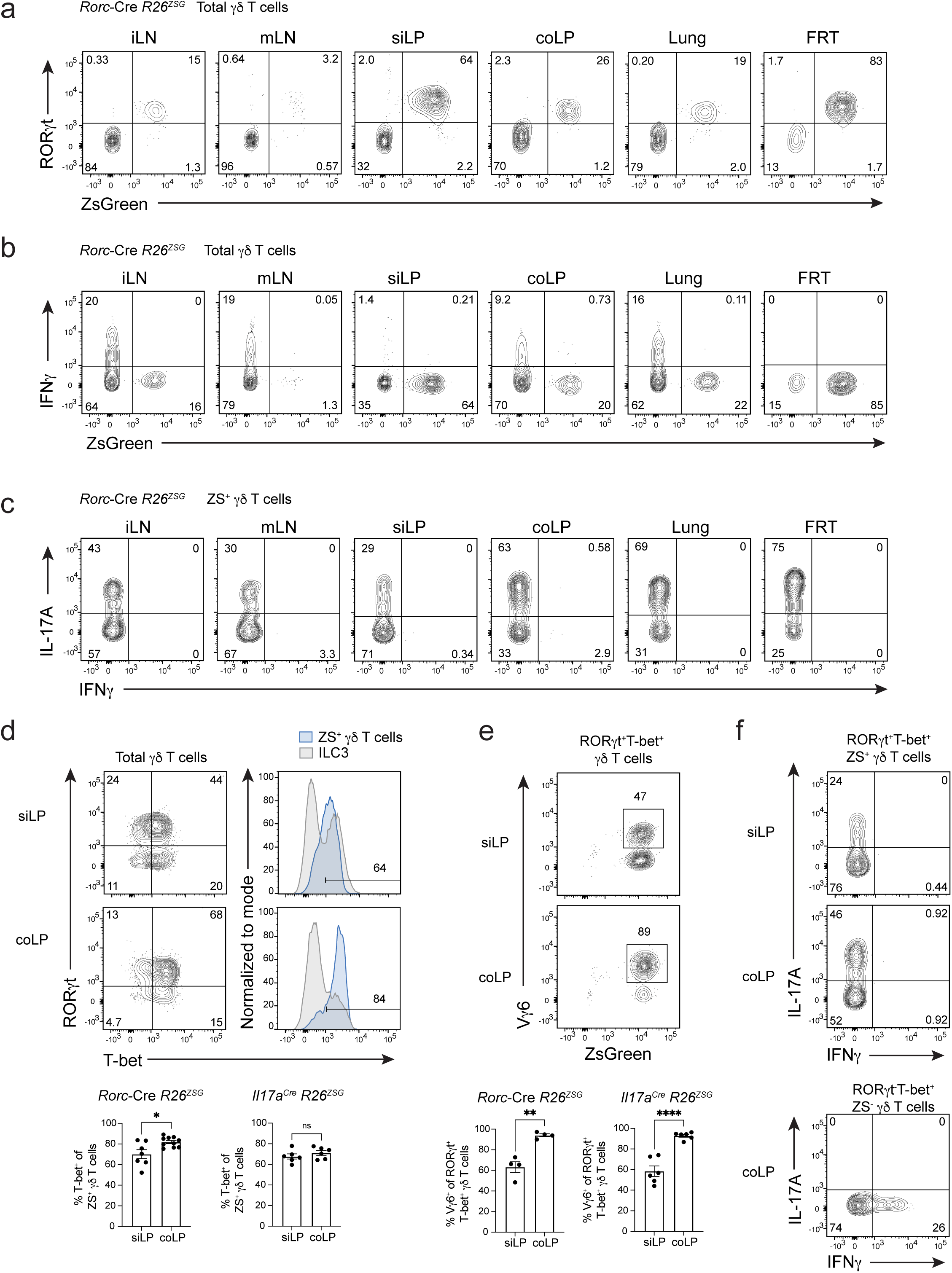
Tγδ17 cell identity is stable at steady state and IFNγ production is restrained in intestinal T-bet+ Tγδ17 cells. (**a**) Flow cytometric analysis of RORγt versus ZS-green expression in total γδ T cells (CD3^+^γδTCR^+^TCRβ^−^) from *Rorc*-Cre *R26^ZSG^*mice. Flow cytometric analysis of cytokine production by (**b**) total γδ T cells (CD3^+^γδTCR^+^TCRβ^−^) and (**c**) ZS^+^ CD3^+^γδTCR^+^TCRβ^−^ cells from lymphoid (iLN and mLN) and nonlymphoid tissues (siLP, coLP, Lung, FRT) of *Rorc*-Cre *R26^ZSG^* mice after 4 hours of PMA/Ionomycin stimulation. Representative of three or more independent experiments with n=8 or more, except lung n=5. (**d**) Flow cytometric analysis of RORγt and T-bet expression in siLP and coLP γδ T cells (CD3^+^γδTCR^+^TCRβ^−^) from *Rorc*-Cre *R26^ZSG^* mice. ILC3 provided as T-bet^+^ and T-bet^−^ reference. (**e**) As in (a), except analysis of Vγ6 and ZS-green expression among RORγt^+^T-bet^+^ γδ T cells. (**f**) Cytokine production after PMA/Ionomycin stimulation for 4 hours of siLP and coLP RORγt^+^T-bet^+^ ZS^+^ γδ T cells (top) and coLP RORγt^−^T-bet^+^ ZS^−^ γδ T cells (bottom). Three or more independent experiments performed. **d, e,** and **f** summary plots are from two independent experiments; (**d**) left n=7-10, (**d**) right n=6, (**e**) left n=4, (**e**) right n=6 mice. All results represent mean ± s.e.m. *P < 0.05; **P < 0.01; ****P < 0.0001; ns, not significant (two-tailed unpaired Student’s t-test). Numbers in flow plots represent percentages of cells in the gate.

Tissue-resident lymphocytes have distinct transcriptional programming dependent on local tissue-derived signals^3,39^. Of the tissues assayed, the intestine is unique in harboring a robust population of RORγt^+^ T-bet^+^ γδ T cells (Extended Data Fig. 1c), which represent interesting candidates for evaluating plasticity. Intestinal Tγδ17 cells upregulate T-bet downstream of environmental cues in the lamina propria during neonatal life^30^. The majority of intestinal Tγδ17 cells co-expressed T-bet in both *Rorc*-Cre *R26^ZSG^* and *Il17a*^Cre^ *R26^ZSG^* mice (Fig.1d). Among T-bet^+^ Tγδ17 cells, 60% and 90% were Vγ6^+^ in the siLP and coLP, respectively (Fig. 1e), as previously reported^30^. In naive mice, intestinal T-bet^+^ Tγδ17 cells failed to produce IFNγ after ex vivo stimulation, despite expressing high levels of T-bet (Fig. 1f). Thus, T-bet expression in Tγδ17 cells is not sufficient for IFNγ production during homeostasis, suggesting that steady state intestinal Tγδ17 cells are poised for type 1 plasticity, and additional signals are required for IFNγ production.

### Vγ6^+^ Tγδ17 cells are functionally plastic after intestinal *S. typhimurium* infection

To test whether a type 1 intestinal infection could elicit effector conversion of T-bet^+^ Tγδ17 cells, we orally infected *Il17a*^Cre^ *R26^ZSG^* or *Rorc*-Cre *R26^ZSG^* fate-mapping mice with *Salmonella typhimurium* and analyzed coLP γδ T cells after 96 hours. *S. typhimurium* infection did induce robust IFNγ production by type 3 fate-mapped ZS^+^ coLP γδ T cells and a modest, but significant, reduction in IL-17A production (Fig. 2a and Extended Data Fig. 1d). As expected, IFNγ-producing ZS^−^ γδ T cells also increased in frequency (3-fold) after infection compared to naive controls (Fig. 2b and Extended Data Fig. 1e). Notably, within the type 3 fate-mapped Tγδ17 cell compartment, Vγ4^+^ and Vγ6^+^ cells had divergent responses (Fig. 2b and Extended Data Fig. 1f, g). Vγ4^+^ Tγδ17 cells maintained high levels of IL-17A production and were less than 1.5% IFNγ^+^ post-infection, consistent with a stable type 3 program (Fig. 2b and Extended Data Fig.1g). In contrast, Vγ6^+^ Tγδ17 cells underwent significant conversion to type 1 effectors, with 25% of ZS^+^ cells producing IFNγ without IL-17A (Fig. 2b). Furthermore, while 30% of Vγ6^+^ Tγδ17 cells maintained IL-17A production after *S. typhimurium* infection, there was a significant decrease in the frequency and level of IL-17A compared to naïve controls (Fig. 2b and Extended Data Fig.1f). Interestingly, the percentage of IL-17A^+^IFNγ^+^ Vγ6^+^ Tγδ17 cells post-infection was very low and the majority of Vγ6^+^ Tγδ17 cells produced either IL-17A or IFNγ (Fig. 2b). This is unlike Th17 cells that generate a prominent transitional IL-17A^+^IFNγ^+^ population during type 1 conversion^17,22^, suggesting that Vγ6^+^ Tγδ17 cells undergo type 1 plasticity with initial full downregulation of IL-17A or from IL-17A-non-expressors. Additionally, the frequency of Vγ6^+^ Tγδ17 cells greatly increased in the mLN after infection and IFNγ was also induced in these cells, mirroring findings from the coLP (Extended Data Fig.1h, i, j). Therefore, intestinal Vγ6^+^ Tγδ17 cells exhibit type 1 effector plasticity in response to *S. typhimurium* infection.

**Fig. 2.**
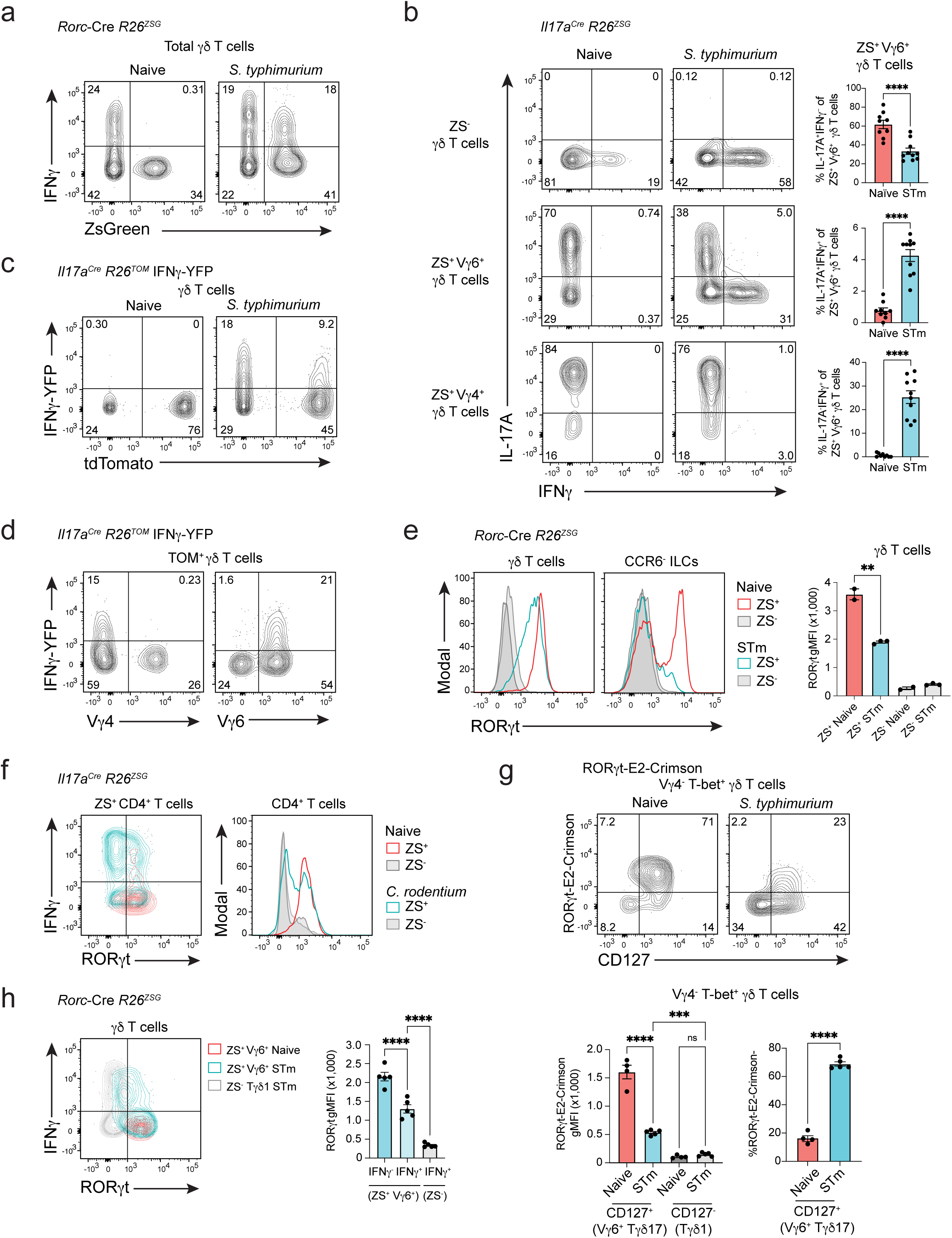
Vγ6^+^ Tγδ17 cells are functionally plastic after intestinal *S. typhimurium* infection. (**a**) Flow cytometric analysis of coLP of naïve and *S. typhimurium* infected *Rorc*-Cre *R26^ZSG^* mice for IFNγ production versus ZS-green expression gated on total γδ T cells. (**b**) Cytokine production from subsets of γδ T cells; ZS^−^ γδ T cells, and ZS^+^ γδ T cells further gated on Vγ4 or Vγ6 from the coLP of *Il17a*^Cre^ *R26^ZSG^* naïve or (STm) *S. typhimurium* infected mice. Summary data pooled from two independent experiments. n=9 or more mice per condition. (**c**) IFNγ-YFP expression in total γδ T cells and (**d**) Vγ4 or Vγ6 versus IFNγ-YFP expression in fate-mapped TOM^+^ γδ T cells from naïve and *S. typhimurium* infected *Il17a*^Cre^ *R26^TOM^*IFNγ-YFP mice. Flow plots representative of more than three independent experiments, n >8 for **c** and **d**. (**e**) RORγt expression among ILCs (CD3^−^CD127^+^CCR6^−^) and γδ T cells for ZS^−^ and ZS^+^ populations from naïve and (STm) *S. typhimurium* infected *Rorc*-Cre *R26^ZSG^*mice. Summary data from one experiment representative of three or more experiments, n >10. (**f**) IFNγ and RORγt expression among CD4^+^ T cells (CD3^+^CD4^+^) for ZS^−^ and ZS^+^ populations from naïve and day 13 *C. rodentium* infected *Il17a*^Cre^ *R26^ZSG^* mice. (**g**) Flow cytometric analysis of coLP of naïve and *S. typhimurium* infected RORγt-E2-Crimson reporter mice for RORγt-E2-Crimson expression gated on CD127^+^ or CD127^−^ cells pregated on Vγ4^−^ T-bet-ZsGreen^+^ γδ T cells. Summary data n= 4-5 mice per condition from one independent experiment for RORγt-E2-Crimson gMFI and frequency. (**h**) Flow cytometric analysis of coLP of naïve and *S. typhimurium* infected *Rorc*-Cre *R26^ZSG^* mice for IFNγ and RORγt expression gated on Vγ6^+^ ZS^+^ γδ T cells and ZS^−^ γδ T cells for comparison. Summary graph from one experiment (n=5) but representative of two or more experiments. Statistical analyses included Two-tailed unpaired Student’s t-tests for **b, e** and ordinary one-way ANOVA test for **g, h**. Numbers in flow plots represent percentages of cells in the gate. All results represent mean ± s.e.m. *P < 0.05; **P < 0.01; ****P < 0.0001; ns, not significant.

To assess IFNγ production directly, we performed an oral *S. typhimurium* infection in *Il17a*^Cre^ *R26^tdTomato(TOM)^* fate-mapping mice bred to an IFNγ-YFP reporter^40^ and analyzed the coLP. This model recapitulated the findings after ex vivo stimulation; *Il17a* fate-mapped TOM^+^ γδ T cells induced IFNγ (YFP^+^) expression and TOM^+^YFP^+^ cells were uniformly Vγ6^+^ (Fig. 2c, d). These results suggest that intestinal Vγ6^+^ and Vγ4^+^ Tγδ17 cells have distinct potentials for plasticity—at least in response to *S. typhimurium* infection—that may reflect their unique ontogenies^11^, TCR specificities, or transcriptional regulation^5^.

Fate-mapping with the *Rorc*-Cre *R26^ZSG^* model allowed us to directly compare Tγδ17 and ILC3 plasticity after intestinal infection. Colonic ILC3 conversion to ILC1s (ex-ILC3s) was marked by a complete downregulation of RORγt expression and was observed in both naïve and *S. typhimurium*-infected mice, as ILC3 plasticity occurs constitutively at steady state (Fig. 2e)^18^. RORγt expression was also fully lost by Th17 cells after plastic conversion to IFNγ^+^ ex-Th17 cells during intestinal *Citrobacter rodentium* infection (Fig. 2f). In contrast, fate-mapped Tγδ17 cells (ZS^+^) only partially downregulated RORγt protein after *S. typhimurium* infection and could be distinguished from bona fide Tγδ1 cells (ZS^−^) based on retained RORγt expression (Fig. 2e and Extended Data Fig. 1k). However, using an RORγt-E2-Crimson reporter strain^41^, we observed a 3-fold reduction in RORγt-E2-Crimson levels post-infection, with a substantial proportion of Vγ6^+^ Tγδ17 cells losing RORγt-E2-Crimson expression (Fig. 2g), indicative of a more substantial downregulation of RORγt transcription compared to protein by 96h. Moreover, IFNγ-producing Vγ6^+^ Tγδ17 cells had significantly reduced RORγt expression compared to IFNγ^−^ counterparts, suggesting that partial RORγt downregulation may be important for type 1 plasticity (Fig. 2h). Together, these results indicate that Tγδ17 cell plasticity takes place after *S. typhimurium* infection, exclusively for the Vγ6^+^ subset, permitting type 1 cytokine production while maintaining intermediate type 3 LDTF expression.

### Single cell multiome characterization of V**γ**6^+^ T**γδ**17 plasticity

To explore the molecular mechanisms governing the stability and plasticity of Tγδ17 cells, we took advantage of the type 1 effector conversion observed for colonic Tγδ17 cells after oral *S. typhimurium* infection. For this, we sorted γδ T cells from 96 h *S. typhimurium*-infected and mock-infected (‘naïve’) *Rorc*-Cre *R26^ZSG^*mice and performed 10x Genomics single cell multiome analysis on the isolated nuclei, whereby chromatin accessibility (scATAC-seq) and gene expression (scRNA-seq) measurements are performed in the same cell. After merging of the naïve and *S. typhimurium* infected datasets, weighted nearest neighbor (WNN) analysis used both RNA-seq and ATAC-seq modalities for dimensionality reduction and UMAP projection^42^. Further cell clustering divided the 2,881 γδ T cells into 14 clusters (C), which segregated broadly based on Tγδ1 versus Tγδ17 effector lineage (Fig. 3a). Most clusters consisted of cells predominantly from either the naïve or *S. typhimurium* condition (Fig. 3b and Extended Data Fig. 2a). Tγδ17 clusters (C0, C1, C6, C7, C8, C9, C12, C13) were identified based on the expression of *Rorc*, *Maf*, *Il23r*, *Il1r1*, and *Il17a,* while Tγδ1 clusters (C2, C3, C4, C5, C10, C11) lacked expression of these defining type 3 genes, and instead expressed key type 1 genes, such as *Il12rb2*, *Tbx21*, *Ifng*, *Gzmb*, and *Tnf* (Fig. 3c and Extended Data Fig. 2b, c). Notably, Tγδ17 clusters had expression of both *Rorc* and *Tbx21* (Fig. 3c), consistent with their protein co-expression (Fig. 1d).

**Fig. 3.**
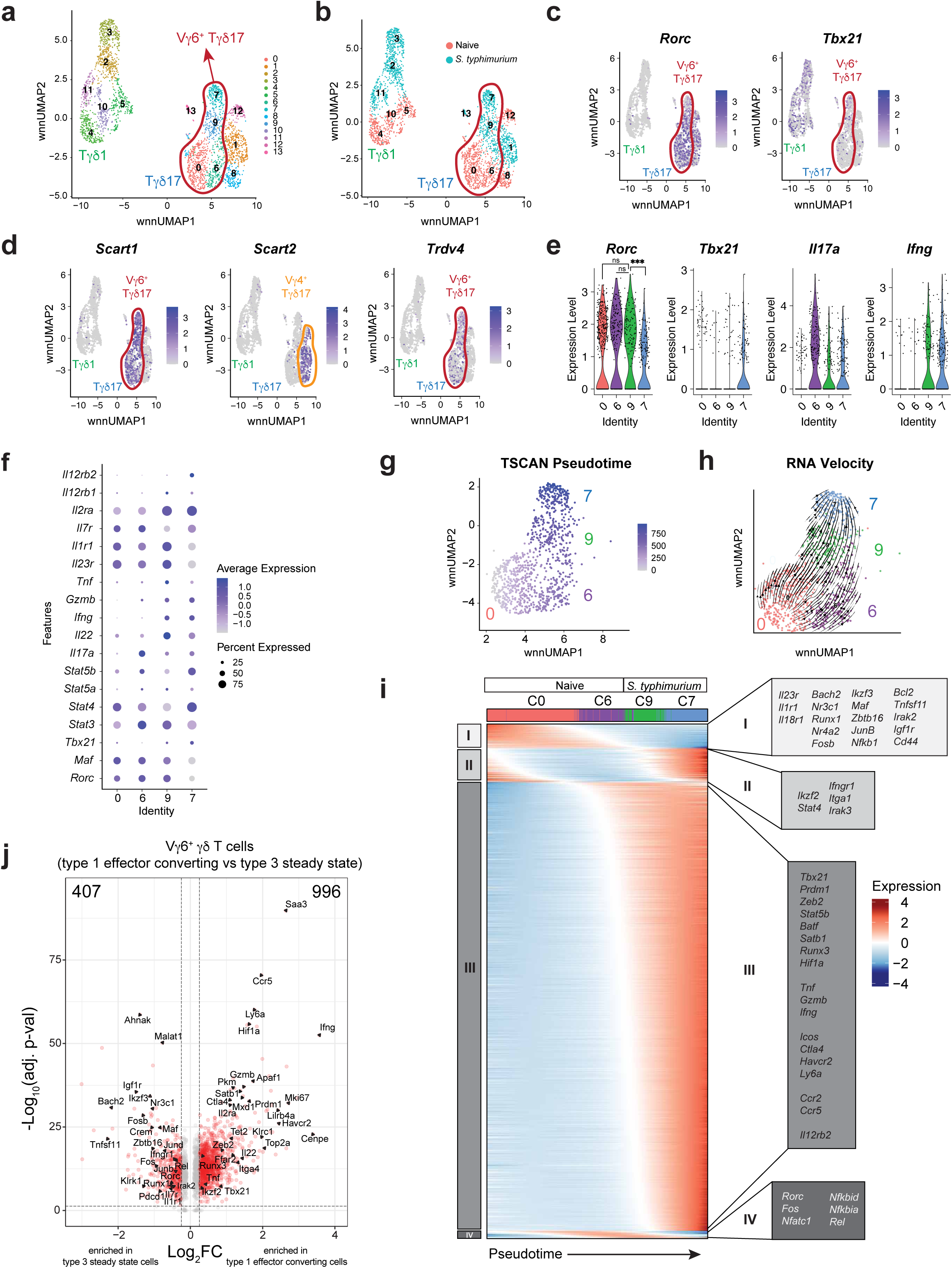
Single cell multiome characterization of Vγ6^+^ Tγδ17 plasticity. (**a**) DimPlot of γδ T cells reclustered with WNN reduction. (**b**) DimPlot of γδ T cells from each condition. (**c**) FeaturePlot showing *Rorc* and *Tbx21* expression in γδ T cells clusters. (**d**) FeaturePlot showing *Cd163l1* (Scart1), *5830411N06Rik* (Scart2), and *Trdv4* expression in γδ T cells clusters. (**e**) Violin Plots of *Rorc*, *Tbx21*, *Il17a*, and *Ifng* expression for Vγ6^+^ Tγδ17 cell clusters. (**f**) DotPlot for select genes in Vγ6^+^ Tγδ17 cell clusters. (**g**) TSCAN pseudotime trajectory on a FeaturePlot of Vγ6^+^ Tγδ17 clusters C0, C6, C9 and C7. (**h**) RNA Velocity (bottom) of Vγ6^+^ Tγδ17 clusters C0, C6, C9 and C7. (**i**) Groups of genes changing along TSCAN pseudotime as in (**g**). (**j**) Volcano plot of post-infection Vγ6^+^ Tγδ17 cell clusters (C7+C9) compared to steady state clusters (C0+C6) with red dots having p-val adj < 0.05 and log2FC > 0.25. One-way ANOVA Tukey test for **e** *Rorc* plot. \**P* < 0.05; \*\**P* < 0.01; ***P < 0.001; \*\*\*\**P* < 0.0001; ns, not significant.

We used the subset-specific transcripts *Scart1* (*Cd163l1*) and *Scart2* (*5830411N06Rik*) to delineate Vγ6^+^ and Vγ4^+^ Tγδ17 cells, respectively^43^ (Fig. 3d). Together with *Trdv4,* which encodes the Vδ1 chain that pairs with the Vγ6 chain on Tγδ17 cells^5^, we identified C0, C6, C7, and C9 as Vγ6^+^ Tγδ17 cells, and C1 and C8 as Vγ4^+^ Tγδ17 cells, with small mixed C12 and C13 (Fig. 3d). C7 and C9 were mostly Vγ6^+^ Tγδ17 cells after *S. typhimurium* infection, whereas C0 and C6 represented mainly steady state Vγ6^+^ Tγδ17 cells (Fig. 3b, d). Interestingly, while steady state cells in C0 and C6 are transcriptionally similar, they differed most significantly in *Il17a* expression, with few low level expressors in C0 and abundant high-level expressors in C6 (Extended Data Fig. 2d and Fig. 3e, f). Thus, steady state Vγ6^+^ Tγδ17 cells comprise cells in either a resting (C0) or a more activated *Il17a*-producing (C6) state, the latter of which also express significantly higher levels of activation-associated *Hif1a*^44^ and *Zeb2*^45^ (Extended Data Fig. 2d). Notably, very little *Trdv4* or type 3 fate-mapped *Zfp506* (ZsGreen) signal was detected among post-infection “Tγδ1” clusters, neither of which was focused in any particular cluster, indicating that Vγ6^+^ Tγδ17 cell plasticity can be characterized within “Vγ6^+^ Tγδ17” clusters (Fig. 3d and Extended Data Fig. 2e). Indeed, with infection-induced plasticity, C9 maintained *Rorc* levels equivalent to steady state C0 and C6 Tγδ17 cells, whereas C7 had significantly decreased *Rorc* expression (Fig. 3e, f). C7 had a negative RNA velocity^46,47^ for *Rorc*, with the lowest levels of both unspliced and spliced transcripts compared to other Vγ6^+^ clusters, consistent with transcriptional downregulation of *Rorc* (Extended Data Fig. 2f). Accordingly, C7 cells also had reduced *Il23r* expression and concomitantly increased *Il12rb2* expression relative to C9 cells, consistent with a type 3 to type 1 shift in identity (Fig. 3f and Extended Data Fig. 2b, c). Similarly, both C9 and C7 cells attenuated *Il17a* and upregulated *Ifng*, *Gzmb*, and *Tnf* expression relative to naïve C6 (Fig. 3f and Extended Data Fig. 2b, c). These graded trends in key lineage-defining genes suggest a conversion of type 3 Vγ6^+^ Tγδ17 cells (C0 and C6) into type 1 effector converted “ex-Tγδ17s” (C7) via a “transitional” C9 following *S. typhimurium* infection. To validate the cellular state transitions among the Vγ6^+^ Tγδ17 cell clusters, we performed RNA velocity analysis^46,47^, and inferred pseudotime trajectories using TSCAN and Monocle 3^48,49^. Together, these analyses corroborated a trajectory for Vγ6^+^ Tγδ17 cell plasticity from steady state (C0 → C6) → transitional C9 → terminal ex-Tγδ17 C7 (Fig. 3g, h and Extended Data Fig. 2g).

We next evaluated the transcriptional changes that occur as Vγ6^+^ Tγδ17 cells undergo effector conversion. For this, genes with significant temporal differences were grouped based on expression patterns along the pseudotime trajectory (Fig. 3i). Interestingly, there was an early downregulation in expression of type 3-stabilizing cytokine receptors (*Il23r*, *Il1r1*, *Il18r1*) and multiple TFs (*Bach2, Nr3c1, Runx1, Nr4a2, Fosb, Ikzf3, Maf, Zbtb16, Junb, Nfkb1*) in Group I that precede the upregulation of Group III genes (Fig. 3i). Many notable genes in Group III, such as TFs (*Tbx21, Prdm1, Zeb2, Stat5b, Batf, Satb1, Runx3, Hif1a*), cytokines (*Tnf, Gzmb, Ifng*), and receptors (*Icos, Ctla4, Havcr2, Ly6a, Ccr2, Ccr5, Il12rb2*), indicate a shift towards type 1 programming and a heightened effector state after infection (Fig. 3i). In line with this, C0 and C6 were enriched for a naïve CD8^+^ T cell gene expression signature, whereas C9 and C7 were enriched in an effector CD8^+^ T cell signature (Extended Data Fig. 2h). Of note, *Rorc* RNA was downregulated late in the trajectory in C7 (Group IV), suggesting that key TFs downregulated in Group I or upregulated in Group III contribute to the eventual attenuation of *Rorc* expression (Fig. 3i). Thus, ex-Tγδ17 cells have undergone a significant global shift in type 3 to type 1 effector programming (Fig. 3j), although, interestingly, they have retained low level *Rorc* expression and their overall transcriptional and chromatin state remains more similar to Tγδ17 than Tγδ1 cells based on the Euclidean distance between clusters in the WNN UMAP (Extended Data Fig. 2i). Taken together, Vγ6^+^ Tγδ17 cell plasticity involves the downregulation of key type 3 genes, followed by a substantial upregulation of type 1 effector genes, representing a functional conversion into ex-Tγδ17 cells.

### TIM-3 marks ex-T**γδ**17 cells with type 1 functionality

To characterize distinguishing features of ex-Tγδ17 cells, we performed differential expression (DE) analysis comparing type 1-converted C7 to transitional C9 Vγ6^+^ γδ T cells (Fig. 4a). Interestingly, *Havcr2,* which encodes co-inhibitory receptor TIM-3, was significantly upregulated in ex-Tγδ17 C7 cells (Fig. 4a). Moreover, *Havcr2* expression was exclusive to C7 and was not expressed by other Vγ6^+^ clusters (C0, C6, or C9), suggesting TIM-3 as a selective marker for Vγ6^+^ ex-Tγδ17 cells (Fig. 4b). Although extensively studied in CD8^+^ T cells, co-inhibitory receptors are also expressed by several tissue-resident lymphocytes at steady state^50^. For example, PD-1 marks Tγδ17 cells during homeostasis^32,43,51^, and restricts IL-17A production by lung and colonic Tγδ17 cells^51,52^. Unlike *Havcr2* expression, the expression of *Pdcd1* (PD-1), *Ctla4*, and co-stimulatory *Icos* was not exclusive to any specific Vγ6^+^ γδ T cell cluster (Fig. 4b). At the protein level, we validated that cell surface TIM-3 expression was selectively induced by type 3 fate-mapped Vγ6^+^ γδ T cells following *S. typhimurium* infection and was not appreciably expressed by either naïve or Vγ4^+^ Tγδ17 cells (Fig. 4c). Consistent with the RNA trends, TIM-3^+^ *Il17a*-fate-mapped Vγ6^+^ γδ T cells from *S. typhimurium*-infected colons were significantly enriched for ex-Tγδ17 cells, producing IL-17A at lower levels and IFNγ at a higher frequency than TIM-3^−^ counterparts after ex vivo stimulation (Fig. 4d). The enhanced capacity of fate-mapped coLP TIM-3^+^ Vγ6^+^ γδ T cells to produce IFNγ was also observed in infected *Il17a*^Cre^ *R26^TOM^* IFNγ-YFP reporter mice, independent of exogenous stimulation (Extended Data Fig. 3a). Interestingly, the TIM-3^+^ Vγ6^+^ γδ T cells also co-expressed PD-1, although at levels significantly lower compared to TIM-3^−^ cells (Extended Data Fig. 3b). This reduced PD-1 expression may facilitate IFNγ expression and proliferation by ex-Tγδ17 cells^53^. Indeed, anti-PD-1 blockade of lung Vγ6^+^ Tγδ17 cells induced cell expansion^51^. Therefore, during *S. typhimurium-*induced plasticity, TIM-3 marks Vγ6^+^ ex-Tγδ17 cells that have enhanced type 1 effector function.

**Fig. 4.**
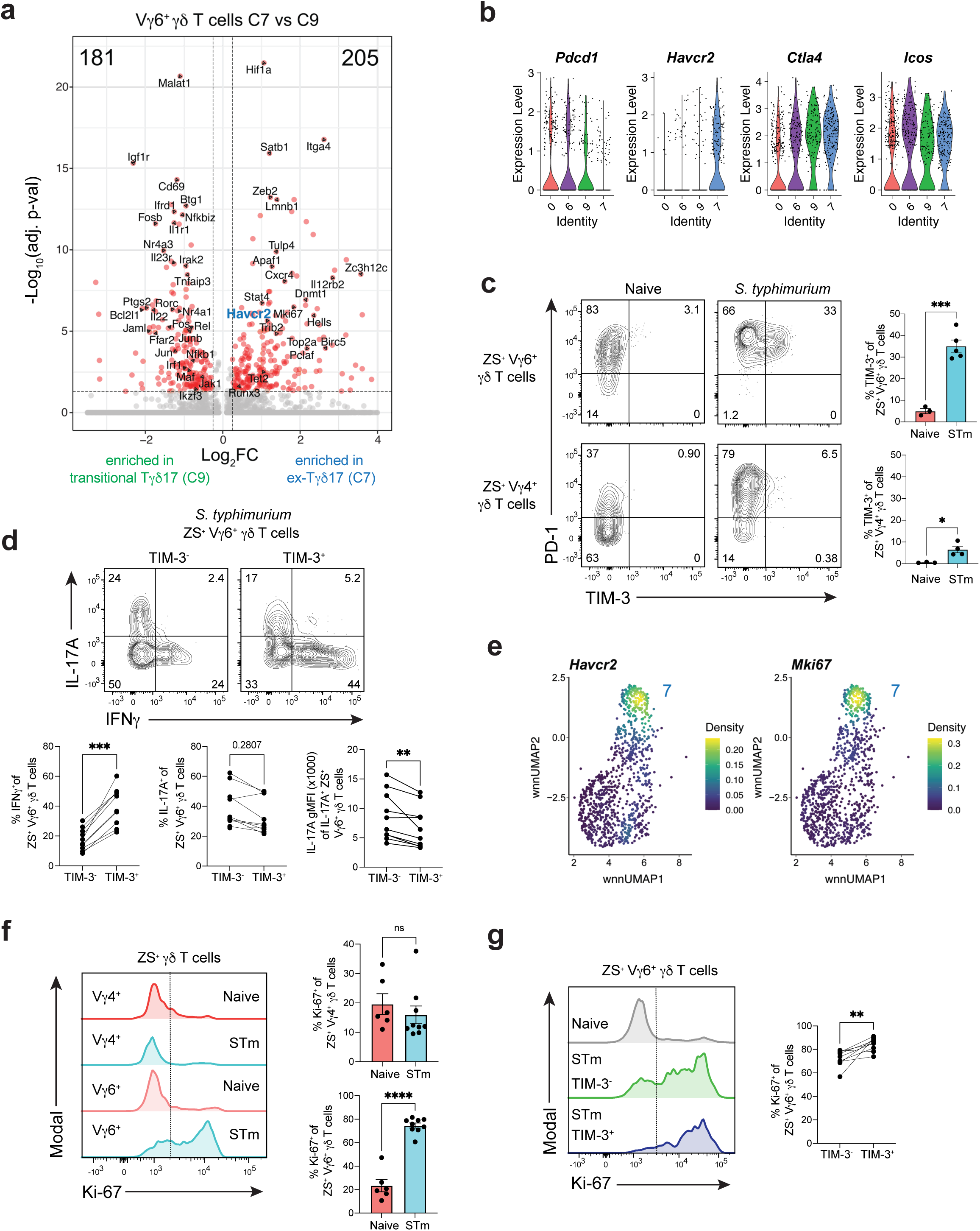
TIM-3 marks ex-Tγδ17 cells with type 1 functionality. (**a**) Volcano plot for differentially expressed genes between C7 versus C9 Vγ6^+^ Tγδ17 cells. Red dots denote significant differences with p-val adj < 0.05 and log_2_FC > 0.25; FC, fold change. (**b**) Violin plots showing expression of select genes in Vγ6^+^ Tγδ17 cell clusters. (**c**) Flow cytometric analysis of PD-1 and TIM-3 expression on ZS^+^ Tγδ17 cells from naive and *S. Typhimurium* (STm) infected *Il17a*^Cre^ *R26^ZSG^*mice. (Top) Vγ6^+^ Tγδ17 cells and (bottom) are Vγ4^+^ Tγδ17 cells gated as Vγ6^−^ Tγδ17 cells. Summary data from one experiment, representative of more than three experiments. (**d**) Flow cytometry plots showing cytokine production after ex vivo stimulation gated on TIM-3^−^ versus TIM-3^+^ Vγ6^+^ Tγδ17 cells from *S. Typhimurium* (STm) *Il17a*^Cre^ *R26^ZSG^*infected mice. Summary plots from two independent experiments n=10 mice. (**e**) Nebulosa density plot for *Havcr2* and *Mki67* for Vγ6^+^ Tγδ17 cell clusters. (**f**) Ki-67 frequency in ZS^+^ Tγδ17 cells from naive and *S. Typhimurium* (STm) infected *Il17a*^Cre^ *R26^ZSG^* or *Rorc*-Cre *R26^ZSG^* mice. Representative histograms gated on Vγ6^+^ or Vγ4^+^ Tγδ17 cells. Summary data compiled from two independent experiments and n=6 naive mice and n=9 STm mice. (**g**) Histogram gated on TIM-3^−^ versus TIM-3^+^ Vγ6^+^ Tγδ17 cells or Vγ6^+^ Tγδ17 cells from a naive mouse as a control. Summary data of percent Ki-67^+^ among TIM-3^−^ versus TIM-3^+^ Vγ6^+^ Tγδ17 cells from *S. Typhimurium* (STm) infected *Il17a*^Cre^ *R26^ZSG^* or *Rorc*-Cre *R26^ZSG^* mice, compiled from two independent experiments and n=9 mice. Statistical analyses included a Two-tailed unpaired Student’s t-test for **c**, **f** and Two-tailed paired Student’s t-test for **d, g**. Results represent mean ± s.e.m. \**P* < 0.05; \*\**P* < 0.01; ***P < 0.001; \*\*\*\**P* < 0.0001; ns, not significant. Numbers in flow plots represent percentages of cells in the gate.

DE analysis revealed numerous genes upregulated in ex-Tγδ17 C7 compared to transitional C9 that are associated with the cell cycle, proliferation, DNA replication, and DNA repair (e.g. *Mki67*, *Top2a*, *Birc5*, *Dnmt1*, *Cdk6*, *Topbp1*, *Atad5*, *Brca1*, *Lmnb1*, *Hells*, *Ncapg*, *Smc2*, *Mcm6*, and *Cenpe*) (Extended Data Fig. 3a). In addition, ex-Tγδ17 C7 had a more apoptotic profile consistent with terminal cells, with significant upregulation of pro-apoptotic *Apaf1* and downregulation of anti-apoptotic *Bcl2l1* and *Tnfaip3* (Extended Data Fig. 3c). Interestingly, *Mki67* (encodes Ki-67), a marker of proliferation, coincided with *Havcr2* expression and was most highly expressed in ex-Tγδ17 C7 (Fig. 4e). By flow cytometry, both Vγ4^+^ and Vγ6^+^ steady state Tγδ17 cells expressed low levels of Ki-67, approximately 20%, consistent with other tissue-resident lymphocytes^54,55^ (Fig. 4f). However, after *S. typhimurium* infection, there was a significant and selective increase in the frequency of ZS^+^ Vγ6^+^ γδ T cells expressing Ki-67, indicative of a proliferative response unique to this subset (Fig. 4f). Notably, a higher frequency of TIM-3^+^ Vγ6^+^ γδ T cells were Ki-67^+^ versus TIM-3^−^ counterparts (Fig. 4g). These results support that, among Tγδ17 cells, the Vγ6^+^ subset is the dominant responder to *S. typhimurium* infection, potentially due to upregulation of the Vγ6Vδ1 TCR ligand in the inflamed environment^56^. Taken together, ex-Tγδ17 cells marked by TIM-3 *in vivo* have enhanced type 1 effector function and a signature of highly proliferative cells, despite co-expressing PD-1 and TIM-3.

### bZIP TFs are dynamically regulated during V**γ**6^+^ T**γδ**17 cell effector plasticity

To uncover the gene regulatory networks (GRNs) controlling Vγ6^+^ Tγδ17 cell effector plasticity, we applied complementary strategies that capitalize on both the RNA expression and chromatin accessibility modalities of the single cell multiome analysis. First, we employed <u>s</u>ingle-<u>c</u>ell r<u>e</u>gulatory <u>n</u>etwork inference and <u>c</u>lustering (SCENIC), a computational method that uses scRNA-seq data to link TFs and their inferred target genes based on co-expression, thus defining ‘regulons’ governing cell state^57^. In line with the trajectory analysis, hierarchical clustering of the Vγ6^+^ γδ T cell regulons revealed greatest similarities between the steady state C0 and C6 populations and transitional C9, with ex-Tγδ17 C7 clustering separately (Fig. 5a). Notably, *Rorc* regulon activity is substantially abrogated in the terminal ex-Tγδ17 C7 (Fig. 5a, group 3), consistent with the late downregulation of *Rorc* mRNA in C7 (Fig. 3e, i). In contrast, while *Tbx21* (T-bet) regulon activity was not detected, the activity of T-bet cofactors, such as Ets1^58^, Runx2^59^, and STAT4^60^ were enhanced in ex-Tγδ17 C7 (Fig. 5a, groups 6, 7), in line with type 1 effector conversion.

**Fig. 5.**
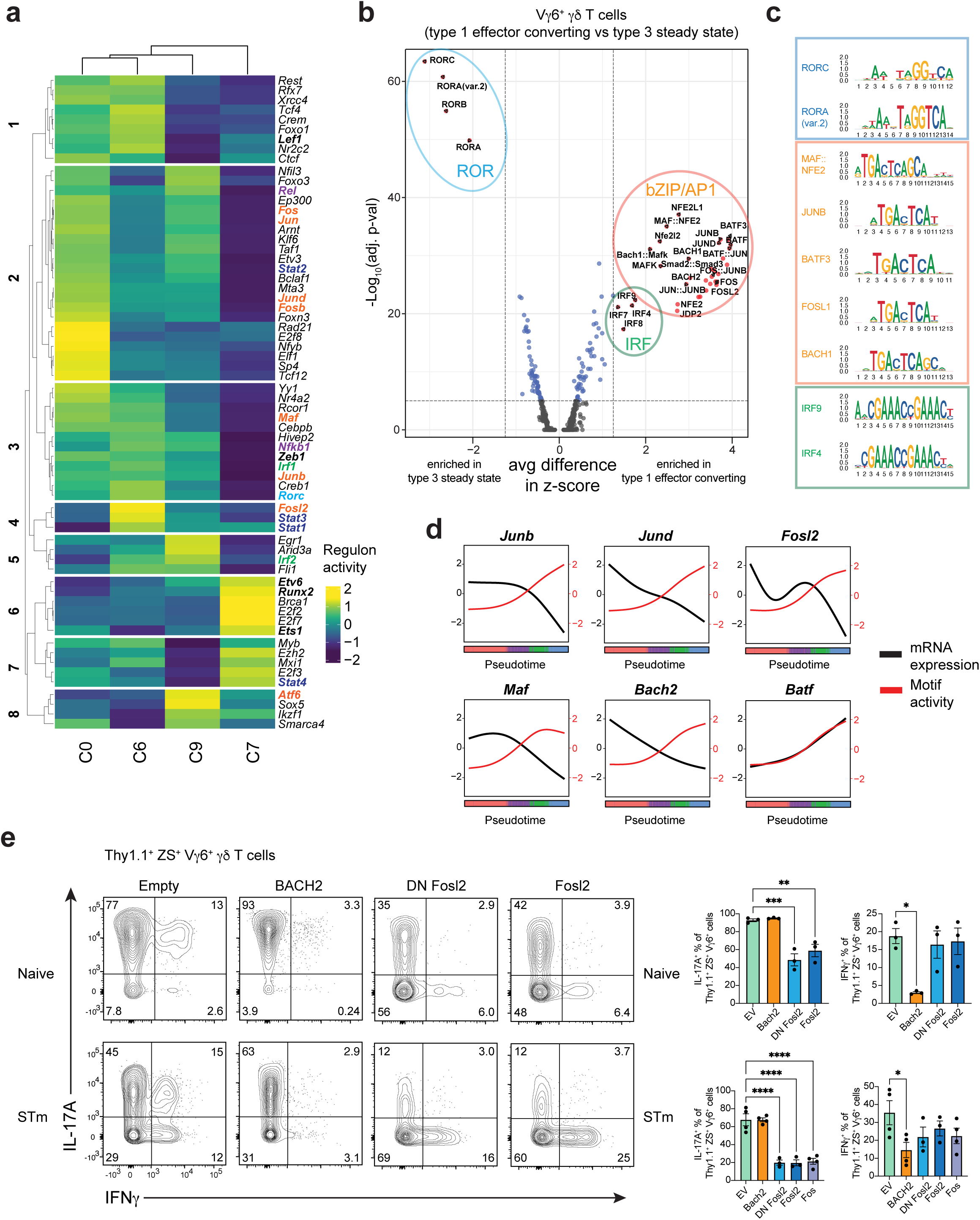
bZIP TFs are dynamically regulated during Vγ6^+^ Tγδ17 cell effector plasticity. (**a**) Regulon activity using SCENIC analysis on Vγ6^+^ Tγδ17 cell clusters C0, C6, C9, and C7. Colored genes draw attention to specific gene families. Red = AP-1 family. (**b**) Motif activity analysis on differentially accessible regions between Vγ6^+^ Tγδ17 cell clusters C7+C9 (post-infection) compared to C0+C6 (steady state). Vertical dotted line represents fold change cut off at 1.25 for average difference in z-score in terms of fold-change between groups. Horizontal line represents p-val adjusted cut off at 5×10^−5^. Colored circles represent TF families. (**c**) Representative top motifs displayed as MotifPlots per TF family circled in (**b**). (**d**) Motif activity (red) and mRNA expression (black) for select TFs across Vγ6^+^ Tγδ17 pseudotime. (**e**) Flow cytometric analysis of cytokine production in 9 day Tγδ17 mLN culture. Gated on transduced Vγ6^+^ (Vγ4^−^) Thy1.1^+^ ZS^+^ γδ T cells after 4h PMA/Ionomycin stimulation. Top row is from naive mLN cultures from *Il17a^Cre^R26^ZSG^* mice and bottom row is from mLN cultures from *S. typhimurium* (STm) infected *Il17a^Cre^R26^ZSG^* mice. Summary graph pooled from two independent experiments for both top and bottom. Statistical analysis includes an Ordinary one-way ANOVA test for **e**. Results represent mean ± s.e.m. \**P* < 0.05; \*\**P* < 0.01; ***P < 0.001; \*\*\*\**P* < 0.0001; ns, not significant. Numbers in flow plots represent percentages of cells in the gate.

SCENIC analysis identified significant changes in activity for several regulons associated with signal-dependent TFs. These TFs connect environmental signals through surface receptors— such as TCRs and cytokine, co-inhibitory, co-stimulatory, and NK receptors—to downstream GRNs controlling γδ T cell identity^8,61^. For example, *Nfkb1* and *Rel* regulon activities were highly diminished in terminal ex-Tγδ17 C7, suggesting a role for decreased NFκB signaling during Vγ6^+^ Tγδ17 cell effector plasticity (Fig. 5a, groups 3, 2). In line with this, the expression of *Il1r1* and *Tnfsf11 (*RANKL)—receptors that activate NFκB signaling and regulate effector identity in type 3 lymphocytes—showed a similar pattern with high levels in steady state Vγ6^+^ Tγδ17 clusters (C0 and C6) that decreased along the trajectory of plasticity (Fig. 3i). Notably, *Stat3 and Stat4* activity were also altered during conversion. In Tγδ17 cells, STAT3 is activated downstream of IL-7R and IL-23R^62,63^. *Stat3* regulon activity was highest in steady state C6, which actively expresses *Il17a*, and progressively decreased from transitional C9 to ex-Tγδ17 C7 (Fig.5a, group 4). Conversely, high *Stat4* regulon activity was specific to C7 (Fig.5a, group 7). Together, this is indicative of a shift from STAT3- to STAT4-driven regulation during Vγ6^+^ Tγδ17 type 1 effector plasticity. This is reminiscent of a similar STAT toggling with type 1 function in NKp46^+^ ILC3s, whereby short duration IL-23R signaling leads to STAT3 activation and IL-22 production, whereas prolonged IL-23R signaling requires STAT4 and T-bet for *Ifng* locus remodeling and IFNγ production^64^. Lastly, and most striking, there was concerted and graded loss in the regulon activity of multiple AP-1 family TFs, including *Jun*, *Junb*, *Jund*, *Maf*, *Fos*, *Fosb*, and *Fosl2* from steady state Vγ6^+^ Tγδ17 cells through C9 and ex-Tγδ17 C7 (Fig. 5a, groups 2, 3, 4). This is in line with the DE analysis in which the TFs downregulated in Vγ6^+^ Tγδ17 cells C9 and C7 were dominated by bZIP/AP-1 superfamily members, such as *Bach2, Jund, Junb, Maf, Fos, Fosb,* and *Fosl2* (Extended Data Fig. 3d). Notably, the downregulation of AP-1 TFs was specific to Vγ6^+^ Tγδ17 cells, as non-plastic Vγ4^+^ Tγδ17 cells maintained the expression of these TFs in post-infection C1 versus steady state C8 (Extended Data Fig. 3e), suggesting they may be relevant regulators of Vγ6^+^ Tγδ17 cell plasticity.

scATAC-seq revealed many regions with differential accessibility as Vγ6^+^ Tγδ17 cells undergo effector conversion (Extended Data Fig. 4a); for example, the *Rorc* locus showed decreased accessibility at multiple non-coding regions, while the *Ifng* locus regions increased in accessibility (Extended Data Fig. 4b). To identify putative regulators that promote or repress functional plasticity in Vγ6^+^ Tγδ17 cells, we performed motif analysis on the global differentially accessible regions for pre- versus post-conversion Vγ6^+^ γδ T cell clusters. The most significant motif activity enriched in the steady state Vγ6^+^ Tγδ17 C0 and C6 were the ROR response elements (Fig. 5b,c and Extended Data Fig. 4c), revealing a striking loss of accessibility at RORγt binding sites during conversion that is consistent with both *Rorc* expression and regulon downregulation. In contrast, motifs enriched with type 1 conversion in the transitional C9 and ex-Tγδ17 C7 were dominated by bZIP/AP-1 and IRF motifs (Fig. 5b,c). Notably, AP-1 motif enrichment was enhanced in both transitional C9 and ex-Tγδ17 C7, whereas IRF motif activity is selectively enhanced in ex-Tγδ17 C7, implying the successive accessibility of distinct sets of regulatory elements during effector plasticity (Extended Data Fig. 4c). The two most significantly enriched motif groups within the bZIP class were for the NFE2 subfamily, encompassing BACH and NFE2-like members, and for the AP-1 superfamily including Jun, Fos, Maf, and BATF members (Fig. 5c). BACH TFs heterodimerize with small Maf proteins and bind Maf-recognition elements (MAREs)^65^, therefore, BACH, Maf, NFE2-like, and AP-1 consensus motifs are highly similar (Fig. 5c). Comparison of TF expression and motif activity in differentially accessible regions revealed an interesting opposing pattern for several bZIP TFs including *Junb, Jund, Fosl2, Maf,* and *Bach2—*but not *Batf*—whereby TF expression declined with conversion, while motif activity steadily increased along the pseudotime trajectory (Fig. 5d). This implies a dominant repressive function for these TFs, with silenced target chromatin becoming more accessible as the negative regulatory TF expression is attenuated. Thus, motif activity analysis suggests that Vγ6^+^ Tγδ17 cell identity is stabilized by the repressive action of a network of bZIP TFs.

To identify putative plasticity regulators among candidates identified in our analyses, we conducted a TF overexpression screen *in vitro* using Tγδ17 cells expanded from the mLN of naïve and *S. typhimurium* infected *Il17a*^Cre^ *Rosa26*^ZSG^ mice. We retrovirally transduced cells with a TF-expressing or empty retroviral vector on day 5 of culture and evaluated the effector cytokine profiles of transduced Thy1.1^+^ ZS^+^ Vγ6^+^ Tγδ17 cells on day 9. Despite originating from naïve mLNs, Vγ6^+^ Tγδ17 cells produced IFNγ after prolonged (6 days) IL-23 and IL-1β stimulation, as previously observed (Fig. 5e)^28^. However, Vγ6^+^ γδ T cells expanded from the mLN of *S. typhimurium* infected mice had greater proportions of IFNγ^+^ and ex-Tγδ17 cells, thus better facilitating evaluation of the IFNγ repression capacity of candidate TFs (Fig. 5e). In total, we tested 16 TFs implicated in Vγ6^+^ Tγδ17 cell plasticity based on altered expression, regulon activity, or motif activity along the conversion trajectory (see Extended Data Fig. 4d). Of the TFs tested, only BACH2 overexpression significantly attenuated IFNγ production among the transduced Vγ6^+^ Tγδ17 cells in mLN cultures derived from both naive and *S. typhimurium* infected mice (Fig. 5e and Extended Data Fig. 4e, f). Interestingly, BACH2 binds to *Ifng* locus regulatory elements and BACH2 overexpression also reduces IFNγ production in CD8^+^ T cells^66^. Additionally, BACH2 restrains the type 1 program in in vitro cultured Th17 cells^67^. Thus, the capacity for IFNγ production by Vγ6^+^ Tγδ17 cells after infection may be due, in part, to BACH2 downregulation and high BACH2 expression may prevent IFNγ production by Vγ6^+^ Tγδ17 cells during homeostasis.

Overexpression of most TFs had no impact on the IL-17A or IFNγ production of Vγ6^+^ Tγδ17 cells relative to empty vector controls (Extended Data Fig. 4e, f). Only Fos, Fosl2, and dominant negative (DN) Fosl2, having a carboxyl-terminal truncation^68^, significantly reduced the proportion of IL-17A producers among transduced Vγ6^+^ Tγδ17 cells (Fig. 5e and Extended Data Fig. 4e, f). Fos TFs do not homodimerize and, instead preferentially form heterodimers with Jun family TFs^69^. Therefore, overexpression of wild-type or DN Fos family members can sequester Jun TFs and prevent their activity, thereby mimicking Jun factor deficiency^68,69^. Accordingly, the findings with Fos TFs also suggest a role for Jun members in supporting IL-17A production in Vγ6^+^ Tγδ17 cells. Moreover, JunB supports Th17 cell identity while restricting alternative fates during inflammation^70^ and JunB was the top interacting partner for FOSL1 and FOSL2 in human Th17 cells among the JUN family^71^, making it a strong candidate for regulation of the type 1 Vγ6^+^ Tγδ17 cell effector switch. Overall, these results implicate a network of bZIP TFs including BACH2, and Jun (*Junb* or *Jund*) and Fos (Fos, Fosb, Fosl2) family TFs in the regulation of Vγ6^+^ Tγδ17 cell effector plasticity.

### JunB and Fosl2 stabilize V**γ**6^+^ T**γδ**17 cell identity

The single cell multiomic TF network and motif analysis, combined with findings from the TF overexpression screen, suggest the involvement of various bZIP TFs, including JunB, Fosl2, and BACH2 in regulating the effector identity and type 1 conversion of intestinal Vγ6^+^ Tγδ17 cells. Notably, these factors directly repress the expression of type 1 effectors, such as T-bet or IFNγ, in various innate and adaptive lymphocytes^66,67,70,72,73^. To assess a potential role of these candidate TFs in regulating effector conversion of Vγ6^+^ Tγδ17 cells, we bred mice harboring *Junb*^fl^, *Fosl2*^fl^, or *Bach2*^fl^ conditional alleles to the *Il17a*^Cre^ *Rosa26*^ZSG^ strain, thus enabling simultaneous gene deletion in *Il17a*-expressing cells and tracking of type 3 fate-mapped cells post-deletion. To evaluate dose-dependent TF contributions, we compared LDTF and effector cytokine profiles for ZS^+^ fate-mapped Vγ6^+^ γδ T cells in the coLP of wild-type (*TF*^WT^), conditional heterozygous (*TF*^HET^), and conditional knockout (*TF*^KO^) littermates.

Among bZIP TFs tested, deletion of JunB had the most significant impact on colonic Vγ6^+^ Tγδ17 cell identity at steady state. Loss of JunB in fate-mapped Vγ6^+^ Tγδ17 cells in *Junb*^KO^ mice resulted in a significant 1.4-fold increase in T-bet expression compared to *Junb*^WT^ counterparts, while RORγt expression remained unchanged (Fig. 6a). Moreover, stimulation with PMA/ionomycin revealed a significant reduction in the frequency of IL-17A producing ZS^+^ Vγ6^+^ Tγδ17 cells in *Junb*^KO^ versus *JunB*^WT^ mice. T-bet and IL-17A are similarly regulated by JunB in Th17 cells, where JunB functions in maintaining the type 3 program^70,74^. Interestingly, JunB deletion in steady-state Vγ6^+^ Tγδ17 cells did not elicit IFNγ production or induce effector conversion, despite the enhanced levels of T-bet expression (Fig. 6b). Fosl2 is a known heterodimerization partner for JunB in type 3 cells^71^. However, deletion of *Fosl2* in coLP Vγ6^+^ Tγδ17 cells had no significant effect on RORγt, T-bet, IL-17A, or IFNγ expression in *Fosl2*^KO^ mice compared to *Fosl2*^WT^ controls (Fig. 6c, d). Similarly, despite being a well-documented repressor of cytokines in T cells, in particular of IFNγ^66,67,75^, conditional loss of *Bach2* in *Bach2*^KO^ Vγ6^+^ Tγδ17 cells did not alter the frequency of IL-17A or IFNγ producers at steady state (Fig. 6e). Together, these bZIP genetic perturbation experiments reveal a requirement for JunB in maintaining type 3 identity in innate-like coLP Vγ6^+^ Tγδ17 cells by supporting IL-17A expression and restraining levels of T-bet.

**Fig. 6.**
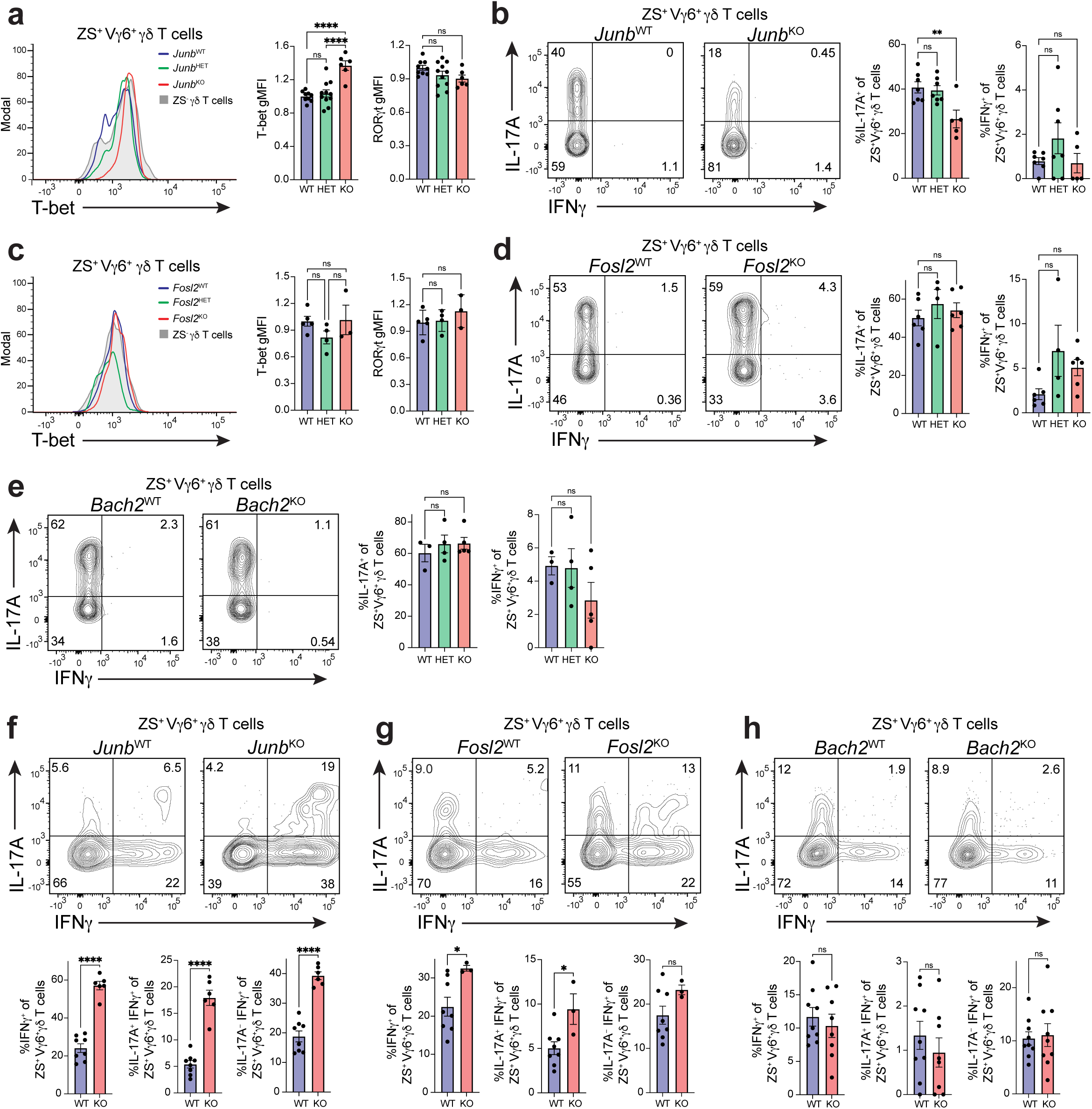
JunB and Fosl2 stabilize Vγ6^+^ Tγδ17 cell identity. (**a-h**) Flow cytometric analysis was performed on colonic ZsGreen^+^ Vγ6^+^ Tγδ17 cells from mice with *Junb*, *Fosl2*, or *Bach2* conditional deletions on the *Il17a^Cre^R26^ZSG^* deleter background at steady state (*TF^+/+^, TF*^WT^; *TF^fl/+^, TF*^HET^; *TF^fl/fl^, TF^KO^*). (**a,c**) Histograms of T-bet expression, and summary plots of normalized T-bet and RORγt expression. (**b,d,e**) Representative flow cytometric analysis and summary plots of the frequency of IL-17A and IFNγ producing cells following 4 h PMA/ionomycin stimulation. (**a-b**) *Junb*: (**a**) *n* = 6–11 mice/genotype; three independent experiments. (**b**) *n* = 5–7 mice/genotype; three independent experiments. (**c-d**) *Fosl2*: (**c**) *n* = 3–5 mice/genotype; two independent experiments. (**d**) *n* = 4–6 mice/genotype; two independent experiments. (**e**) *Bach2*: *n* = 4–5 mice/genotype; two independent experiments. (**f-h**) Representative flow cytometric analysis and summary plots of the frequency of IL-17A and IFNγ producing cells at steady state following 20h stimulation with IL-23 and IL-1β for the following genotypes: (**f**) *Junb* (*n* = 6-8 mice/genotype; two independent experiments), (**g**) *Fosl2* (*n* = 3-8 mice/genotype; two independent experiments), and (**h**) *Bach2* (*n* = 9 mice/genotype; two independent experiments). (**a-e**) Gating was performed on fate-mapped Vγ6^+^ Tγδ17 cells (CD3ε^+^γδTCR^+^TCRβ^−^ZS^+^Vγ6^+^), and for (**f-h**), gating was performed on fate-mapped Vγ4^−^ Tγδ17 cells (CD3ε^+^γδTCR^+^TCRβ^−^ZS^+^Vγ4^−^). Statistical analyses include Ordinary one-way ANOVA tests for (**a-e**) and Two-tailed unpaired Student’s t-tests for (**f-h**). Results represent mean ± s.e.m. \**P* < 0.05; \*\**P* < 0.01; ***P < 0.001; \*\*\*\**P* < 0.0001; ns, not significant. Numbers in flow plots represent percentages of cells in the gate.

Innate-like Tγδ17 cells upregulate IFNγ expression after prolonged IL-23 and IL-1β stimulation^28^. We find that IL-23 and IL-1β also induces type 1 effector plasticity of *ll17a* fate-mapped ZS^+^ coLP Vγ6^+^ Tγδ17 cells, with approximately 10-20% of cells attaining an IL-17A^−^ IFNγ^+^ cytokine converted phenotype after a 20h culture (Fig. 6f-h). Using this *in vitro* plasticity model, we investigated whether loss of JunB, Fosl2, or BACH2 influences type 1 effector conversion of homeostatic coLP Vγ6^+^ Tγδ17 cells. *Il17a* fate-mapped ZS^+^ Vγ6^+^ γδ T cells from *Junb*^KO^ mice had a greater than 2-fold significant increase in the frequency of total IFNγ^+^ producers, IL-17A^+^ IFNγ^+^ double-producers, and IL-17A^−^ IFNγ^+^ cells, resulting in enhanced effector conversion of fate-mapped cells compared to *Junb*^WT^ controls (Fig. 6f). Notably, IL-23 and IL-1β signals revealed a contribution for Fosl2 in suppressing type 1 features as loss of *Fosl2* resulted in a higher frequency of IFNγ production among ZS^+^ Vγ6^+^ γδ T cells (Fig. 6g). In contrast, *Bach2* deficiency had no effect on IFNγ production (Fig. 6h). Thus, among bZIP TFs tested, JunB and Fosl2 function in restricting IFNγ production by Vγ6^+^ γδ T cells during cytokine stimulation-induced effector conversion, while BACH2 is dispensable.

As overnight IL-23 and IL-1β stimulation lowered the threshold to detect a role for JunB and Fosl2 in suppressing IFNγ production, we examined potential redundancy or compensation between these AP-1 TFs in limiting type 1 effector conversion of Vγ6^+^ Tγδ17 cells at steady state. For this, we generated compound conditional *Junb* and *Fosl2* mutants using the *Il17a*^Cre^ *Rosa26^ZSG^* fate-mapping deleter strain. Notably, double conditional deletion of *Junb* and *Fosl2* in Vγ6^+^ Tγδ17 cells was insufficient to result in type 1 conversion or significant IFNγ de-repression following ex vivo stimulation (Extended Data Fig. 5a). This suggests that at steady state, the function of JunB and Fosl2 is redundant with that of other factors that restrict type 1 plasticity or are required for activating conversion downstream of inductive signals. Moreover, reducing the dosage of either JunB or Fosl2 via heterozygosity, or complete double conditional deletion of *Junb* and *Fosl2* in the context of a 20h IL-23 and IL-1β stimulation did not reveal a mutually additive or synergistic role for JunB and Fosl2 in regulation of IFNγ production in fate-mapped Vγ6^+^ γδ T cells (Extended Data Fig. 5b). Specifically, while deletion of *Junb* on a *Fosl2*^KO^ background significantly enhanced the proportion of IFNγ producers and IL-17A^−^ IFNγ^+^ effector converted Tγδ17 cells, deletion of *Fosl2* on a *Junb*^KO^ background had no effect, revealing a more prominent role for JunB in restraining type 1 plasticity in Vγ6^+^ Tγδ17 cells. Thus, the activity of JunB can provide some compensation for loss of Fosl2, whereas Fosl2 cannot compensate for the loss of JunB in steady state Vγ6^+^ Tγδ17 cells stimulated *in vitro* with plasticity-inducing cytokines.

### JunB and Fosl2 sustain ROR**γ**t expression and limit type 1 plasticity of V**γ**6^+^ T**γδ**17 cells during *S. typhimurium* infection

To further evaluate the requirement for bZIP TFs in type 1 plasticity of coLP Vγ6^+^ Tγδ17 cells, we assessed the effector phenotypes of TF conditional mutants during physiological conversion *in vivo*, 96 hours after infection with *S. typhimurium*. Loss of either JunB or Fosl2 resulted in a comparable striking and dose-dependent enhancement in type 1 effector conversion of *Il17a* fate-mapped ZS^+^ Vγ6^+^ Tγδ17 cells (Fig. 7a-d). In particular, there was a marked 1.7- and 1.6-fold reduction in RORγt expression for *Junb*^KO^ and *Fosl2*^KO^ ZS^+^ Vγ6^+^ γδ T cells, respectively, relative to wild-type littermate controls, whereas T-bet levels were unaffected (Fig. 7a, b). This was accompanied by a significant reduction in the proportion of IL-17A producers and a concomitant 2- and 2.4-fold increase in the frequency of IFNγ production by *Junb*^KO^ and *Fosl2*^KO^ ZS^+^ Vγ6^+^ γδ T cells, respectively (Fig. 7c, d). Taken together, these findings are consistent with an enhanced generation of effector-converted Vγ6^+^ ex-Tγδ17 cells in *Junb*^KO^ and *Fosl2*^KO^ versus wild-type control colons during infection, revealing a critical role for both JunB and Fosl2 in restraining type 1 plasticity in Vγ6^+^ Tγδ17 cells in the context of inflammation. Notably, this contrasts with the lack of a similar role for JunB and Fosl2 in suppressing conversion at steady state. The difference may be attributable to the global and concerted loss of AP-1 TF expression and activity in the context of infection that may lower the threshold for detecting a role for individual AP-1 factors and implies that AP-1 TFs may otherwise compensate for one another at steady state.

**Fig. 7.**
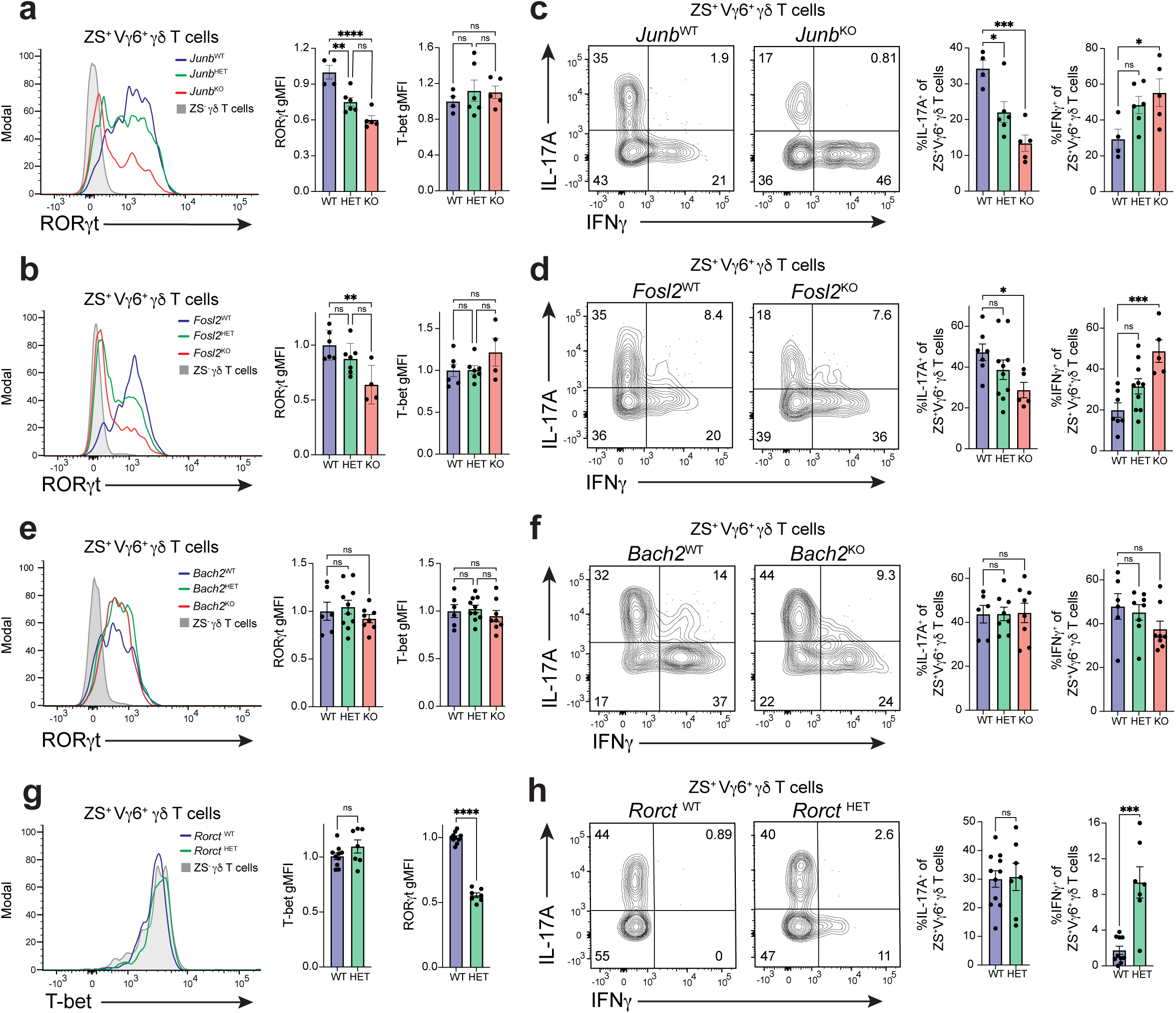
JunB and Fosl2 limit type 1 plasticity in Vγ6^+^ Tγδ17 cells during *S. typhimurium* infection. (**a-f**) Flow cytometric analysis was performed on colonic ZsGreen^+^ Vγ6^+^ Tγδ17 cells from mice with *Junb*, *Fosl2*, or *Bach2* conditional deletions on the *Il17a^Cre^R26^ZSG^* deleter background during *S. typhimurium* infection: (*TF^+/+^, TF*^WT^; *TF^fl/+^, TF*^HET^; *TF^fl/fl^, TF^KO^*). (**a,b,e**) Histograms of RORγt expression, and summary plots of normalized RORγt and T-bet expression. (**c,d,f**) Representative flow cytometric analysis and summary plots of the frequency of IL-17A and IFNγ producing cells following 4 h PMA/ionomycin stimulation. (**a,c**) *Junb*: *n* = 4–6 mice/genotype; two independent experiments. (**b,d**) *Fosl2*: (**b**) *n* = 4–7 mice/genotype; two independent experiments. (**d**) *n* = 6–10 mice/genotype; two independent experiments. (**e-f**) *Bach2*: (**e**) *n* = 6– 10 mice/genotype; two independent experiments. (**f**) *n* = 6–8 mice/genotype; two independent experiments. (**g-h**) Flow cytometric analysis of colonic ZsGreen^+^ Vγ6^+^ Tγδ17 cells from *Rorct^+/+^Il17a^Cre^R26^ZSG^* (*Rorct* ^WT^) and *Rorct^fl/+^Il17a^Cre^R26^ZSG^* (*Rorct* ^HET^) mice at steady state: (**g**) Histograms of T-bet expression, and summary plots of normalized T-bet and RORγt expression (*n* = 7–10 mice/genotype; three independent experiments). (**h**) Representative flow cytometric analysis and summary plots of the frequency of IL-17A and IFNγ producing cells following 4 h PMA/ionomycin stimulation (*n* = 7–10 mice/genotype; three independent experiments). (**a-f**) Gating was performed on fate-mapped Vγ6^+^ Tγδ17 cells (CD3ε^+^γδTCR^+^TCRβ^−^ZS^+^Vγ6^+^), and for (**g-h**), gating was performed on fate-mapped Vγ4^−^ Tγδ17 cells (CD3ε^+^γδTCR^+^TCRβ^−^ZS^+^Vγ4^−^). For *S. typhimurium* infection experiments (**a-f**), cells harvested from mice 96h post-infection. Statistical analyses include Ordinary one-way ANOVA tests for (**a-f**), and a Two-tailed unpaired Student’s t-test for (**g-h**). Results represent mean ± s.e.m. \**P* < 0.05; \*\**P* < 0.01; ***P < 0.001; \*\*\*\**P* < 0.0001; ns, not significant. Numbers in flow plots represent percentages of cells in the gate.

The role of BACH2 in Vγ6^+^ Tγδ17 cell plasticity is less clear. While TF overexpression *in vitro* indicates BACH2 is sufficient to suppress IFNγ expression in Vγ6^+^ Tγδ17 cells (Fig. 5e), *Bach2* deficiency had no effect on Vγ6^+^ Tγδ17 cell effector identity at steady state or on conversion following stimulation with IL-23 and IL-1β (Fig. 6i). Similarly, type 1 plasticity of fate-mapped coLP ZS^+^ Vγ6^+^ Tγδ17 cells induced by *S. typhimurium* infection was unaltered in *Bach2*^KO^ compared to *Bach2*^WT^ mice, with no significant differences in T-bet or RORγt expression, nor in the frequency of IL-17A and IFNγ-producing cells (Fig. 7e, f), confirming that BACH2 is not an obligatory regulator of Vγ6^+^ Tγδ17 cell identity. Interestingly, conflicting outcomes were observed in a mLN Tγδ17 cell expansion culture context *in vitro* in which loss of *Bach2* resulted in a significant increase in the proportion of IFNγ^+^ ZS^+^ Vγ6^+^ γδ T cells at day 9 of culture (Extended Data Fig. 5c), suggesting that BACH2 can contribute to restriction of IFNγ production in some contexts and that there may be other factors that compensate for *Bach2* deficiency in vivo. Taken together with the more pronounced effect of *Junb* and *Fosl2* deficiency on Vγ6^+^ Tγδ17 cell plasticity during inflammatory cytokine stimulation or *S. typhimurium* infection versus steady state, the data suggests that there are multiple redundant mechanisms stabilizing Vγ6^+^ Tγδ17 cell effector identity during homeostasis.

We next explored the mechanism of AP-1-mediated restraint of type 1 plasticity. *Rorc* is an attractive candidate as enhanced effector conversion of Vγ6^+^ Tγδ17 cells following either *Junb* or *Fosl2* deletion was accompanied by significant downregulation of RORγt expression (Fig. 7 a-d). Moreover, RORγt-related motif activity is the most significant and prominent motif lost within differentially accessible chromatin during Vγ6^+^ Tγδ17 cell plasticity (Fig. 5b), suggesting that attenuation of RORγt function is a critical component of effector conversion. Thus, we assessed whether RORγt downregulation alone is sufficient to induce type 1 conversion in Vγ6^+^ Tγδ17 cells. For this, we genetically reduced RORγt levels using conditionally heterozygous *Rorct*^HET^ mice on the *Il17a*^Cre^ *Rosa26*^ZSG^ fate-mapping deleter background. This decreased RORγt expression levels by 1.8-fold in steady state coLP ZS^+^ Vγ6^+^ Tγδ17 cells from *Rorct*^HET^ relative to *Rorct*^WT^ mice (Fig. 7g). Importantly, this reduction was comparable to the physiological 1.7-fold downregulation of RORγt observed in wild-type fate-mapped Vγ6^+^ Tγδ17 cells during *S. typhimurium-*induced plasticity (Extended Data Fig. 1k). Notably, heterozygous levels of RORγt in *Rorct*^HET^ mice resulted in a significant derepression of IFNγ production up to an average of 9.3% of ZS^+^ Vγ6^+^ γδ T cells (Fig. 7g), without altering T-bet expression levels (Fig. 7g), nor the frequency of IL-17A producers when compared to *Rorct*^WT^ mice (Fig. 7h). *Rorct*^HET^ mice were the only genetic perturbation in our study that was sufficient to result in the aberrant generation of ex-Tγδ17 cells in steady state colons, revealing the importance of RORγt in maintaining type 3 effector stability in Vγ6^+^ Tγδ17 cells. Taken together, regulation of RORγt expression levels is a critical node in the AP-1 network controlling Vγ6^+^ Tγδ17 cell effector plasticity.

## Discussion

Pioneering studies identified Tγδ17 cell functional plasticity in various inflammatory states by the co-production of IL-17A and IFNγ^15,29^. However, the potential for effector conversion into Tγδ1-like cells, the physiological features and contexts of type 1 conversion, and the molecular regulators of innate Tγδ17 cell plasticity remain mostly undefined. In this study, we combine type 3 fate-mapping and single cell multiome approaches with genetic perturbations in *Il17a-*expressing γδ T cells to characterize the conditions supporting the stability versus flexibility of Tγδ17 cells, and the GRNs underlying type 1 plasticity. Together, our findings define and validate an AP-1 regulatory axis centered around JunB and Fosl2 that restricts type 1 effector conversion of Tγδ17 cells in inflammatory contexts.

Type 3 fate-mapping characterization of multiple lymphoid and non-lymphoid tissues at steady state confirms the view that Tγδ17 cell effector identity is exceptionally stable^22^. Prior studies have reported the presence of IL-17A^+^ IFNγ^+^ Tγδ17 cells in inflammatory contexts of infection and cancer^15,28,29^. However, without the use of fate-mapping strategies, it was not previously appreciated that Tγδ17 cells undergoing plasticity can downregulate RORγt expression, extinguish IL-17A expression, and undergo functional effector conversion into single IFNγ producers, as observed following intestinal *S. typhimurium* infection. In future studies, it will be important to evaluate fate-mapped Tγδ17 cell plasticity in additional inflammatory models of infection, autoimmunity, and cancer.

Tγδ17 cell plasticity has distinct and overlapping features with type 1 conversion in other type 3 lymphocytes. During *S. typhimurium*-induced plasticity, Vγ6^+^ ex-Tγδ17 cells significantly downregulate *Rorc* transcription yet retain intermediate levels of RORγt protein expression. In contrast, CCR6^−^ ILC3 and Th17 cell type 1 conversion is accompanied by full shutdown of RORγt expression, such that ex-ILC3s and ex-Th17 cells are phenotypically indistinguishable from *bona fide* ILC1s and Th1 cells, respectively^38^. However, plasticity may be functionally similar among these type 3 lymphocytes irrespective of RORγt protein levels, as there is a striking loss of RORγt function based on GRN regulon activity and motif activity within accessible chromatin in ex-Tγδ17 cells. Nevertheless, there are notable context-dependent differences in plasticity between lineages. While innate-like Vγ6^+^ Tγδ17 cells and CCR6^−^ ILC3s both co-express T-bet at homeostasis and are poised for type 1 conversion^30,76^, only ILC3s undergo type 1 conversion at steady state^18^. Rather, Tγδ17 cell plasticity resembles that of adaptive Th17 cells in which select inflammatory settings trigger conversion to type 1 effectors. In particular, Tγδ17 plasticity occurs during intestinal *S. typhimurium* infection, but not in the context of experimental autoimmune encephalomyelitis (EAE); whereas Th17 cell plasticity is induced by EAE, intestinal infections with *Citrobacter rodentium* or *Helicobacter hepaticus*, and *Staphylococcus aureus* sepsis, but not during cutaneous *Candida albicans* infection^17,22,77,78^.

Functional plasticity of lymphocytes is preceded by the co-expression of relevant LDTFs. Nevertheless, co-expression of T-bet and RORγt alone is insufficient to induce IFNγ production in type 3 lymphocytes. In Th17 cells, additional auxiliary TFs, such as Runx3 are required^79^. Although innately programmed intestinal Tγδ17 cells constitutively co-express RORγt and T-bet, we find IFNγ production is restrained at steady state, similar to CCR6^−^ ILC3s^80^, and additional environmental signals are necessary for induction. Interestingly, heterozygous levels of RORγt in *Rorct*^HET^ Vγ6^+^ Tγδ17 cells—matching the downregulation that occurs during infection*-*induced plasticity—is sufficient to release low level IFNγ production at steady state, implicating *Rorc* as a critical regulatory target for type 1 effector conversion. Indeed, loss of either JunB or Fosl2 in Vγ6^+^ Tγδ17 cells during *S. typhimurium* infection significantly attenuates RORγt expression, suggesting the physiological downregulation of these AP-1 TFs during plasticity as a mechanism to both reduce expression and maintain intermediate levels of RORγt in ex-Tγδ17 cells. Thus, for Vγ6^+^ Tγδ17 cells, downregulation of RORγt rather than upregulation of T-bet protein appears to be the critical determinant of type 1 plasticity. As in other type 3 lymphocytes, the balance of these two LDTF determines effector transitions and plasticity^41,80,81^, as RORγt and T-bet compete for transcriptional cofactors^79^.

AP-1 factors are critical regulators of intestinal Vγ6^+^ Tγδ17 cell effector identity and plasticity. We show that type 1 conversion of Vγ6^+^ Tγδ17 cells is characterized by a global and concerted loss of AP-1 TF expression and regulatory activity and identify non-redundant roles for JunB and Fosl2 in restraining type 1 plasticity. Additionally, AP-1 binding sites are among the most highly enriched motifs in regions gaining accessibility in Vγ6^+^ ex-Tγδ17 cells, suggesting that AP-1 TFs control Vγ6^+^ Tγδ17 cell effector flexibility through regulation of the active regulatory landscape. These findings are reminiscent of the requirement for AP-1 TFs in regulating chromatin accessibility and plasticity in Th17 cells^70,72^. Indeed, similarly in Vγ6^+^ Tγδ17 and Th17 cells, JunB represses T-bet, while both JunB and Fosl2 limit type 1 effector conversion during inflammation^70,72^. JunB also selectively supports RORγt expression in the context of inflammation in Vγ6^+^ γδ T cells, as in pathogenic Th17 cells^82^. Thus, although innate-like Tγδ17 cells share critical regulators of plasticity with adaptive Th17 cells, ILC3s employ a distinct AP-1 family class (i.e. c-Maf) to restrict type 1 conversion^80^. This difference potentially reflects shared plasticity-inducing GRNs downstream of TCR and cytokine signals in Tγδ17 and Th17 cells.

In Th17 cells, JunB heterodimerizes with BATF and Fosl2^83,84^, however the AP-1 dimerization partners in Tγδ17 cells are not defined. The parallel Tγδ17 effector phenotypes for *Junb* and *Fosl2* deficiency following *S. typhimurium*-induced plasticity suggest these AP-1 TFs function in the same complex. However, this is at odds with the dominant role of JunB at steady state, whereby JunB is required to maintain type 3 identity by supporting IL-17A and limiting T-bet in Vγ6^+^ Tγδ17 cells, while Fosl2 is dispensable. This discrepancy may be attributable to the described role of JunB in regulating the expression of other AP-1 TFs (i.e. *Jun, Batf*, *Fosl2*, and *Jund* in Th17 cells^70^), thus loss of JunB in steady state Tγδ17 cells may disrupt the nature of AP-1 complexes by broadly perturbing dimerization partners. Additionally, Fosl2 may be redundant with Fos and Fosb for heterodimerization with JunB at steady state but indispensable during infection as Jun and Fos family TFs become more limiting. Interestingly, unlike Jun and Fos family members, expression of *Batf* is upregulated during Vγ6^+^ Tγδ17 plasticity, and the motif activity for IRF4, a cooperative binding partner for JunB-BATF^85^, is selectively enriched in ex-Tγδ17 cells. It is tempting to speculate that BATF and IRF4 may shift the balance of AP-1 complex constituents and function from ones that repress to those that activate chromatin accessibility and gene expression supporting type 1 conversion.

The requirement for JunB and Fosl2 in limiting type 1 conversion of Vγ6^+^ Tγδ17 cells is observed in the context of inflammatory signals such as IL-23 and IL-1β stimulation or *S. typhimurium* infection, but not at steady state. This implies that there may be redundant regulatory mechanisms stabilizing the effector identity of innate Tγδ17 cells during homeostasis. Indeed, other AP-1 TFs are very good candidates for compensatory regulators. During *S. typhimurium* infection, the significant global decrease in Jun and Fos family members may substantially impact the concentration of available AP-1 TF binding partners, resulting in destabilization of AP-1 activating and repressive functions, and allowing for concomitant loss of Vγ6^+^ Tγδ17 cell identity and type 1 conversion. The discrepancy in BACH2-dependency in restraining IFNγ production by Tγδ17 cells in vitro versus being dispensable in vivo also supports the view that there are multiple layers of plasticity restriction in vivo. Although sufficient in repressing IFNγ in other lymphocytes^66^, BACH2 may have a minor contributory role in Tγδ17 cells, with other factors compensating for *Bach2* deficiency in vivo. Lastly, within the network of Tγδ17 cell regulators, miR-146a, which has been shown to limit IFNγ expression and the generation of polyfunctional IL-17A^+^ IFNγ^+^ Tγδ17 cells during *Listeria monocytogenes* infection by targeting *Nod1* expression^31^, may also contribute to combinatorial silencing of plasticity during homeostasis. Importantly, it is also worth considering the alternative scenario in which loss of dominant repression mechanisms—irrespective of compensation—may be insufficient to induce plasticity of Vγ6^+^ Tγδ17 cells at steady state in the absence of inductive signals for type 1 conversion, which remain to be fully characterized.

As major environmental cues affecting Tγδ17 cell cytokine production, IL-23 and IL-1β induce STAT and NF-κB pathways, respectively^51^. Our data implicates both pathways in driving plasticity of colonic Vγ6^+^ Tγδ17 cells. Indeed, type 1 effector conversion required signals that were absent at steady state, yet present during *S. typhimurium* infection. In this regard, overnight culture with IL-23 and IL-1β was sufficient to trigger the generation of IFNγ single-producers from Vγ6^+^ Tγδ17 cells isolated from naive mice. Duration of IL-23 and IL-1β stimulation leads to opposing effector fates for Tγδ17 cells, with short-term exposure resulting in robust IL-17A production, and prolonged stimulation leading to IL-17A^+^IFNγ^+^ co-producers and single IFNγ^+^ producers^28^. Similarly, in NKp46^+^ ILC3s, 4h IL-23 stimulation induces STAT3-dependent IL-22 production, whereas 16h IL-23 stimulation promotes STAT4-dependent IFNγ production and *Ifng* locus remodelling^64^. Notably, the analogous IL-23-responsive distal regions of the *Ifng* locus displayed increased accessibility during type 1 conversion of Vγ6^+^ Tγδ17 cells in the inflammatory context of *S. typhimurium* infection, suggestive of IL-23-mediated derepression of *Ifng*. Accordingly, GRN analysis revealed a shift from *Stat3* to *Stat4* regulon activity along the trajectory of Vγ6^+^ Tγδ17 cell plasticity, coinciding with type 3 to type 1 effector conversion. NF-κB signaling was also altered as evidenced by *Nfkb1* and *Rel* regulon activities that are high in responding steady state Vγ6^+^ Tγδ17 cells and decline with *Il1r1* downregulation during type 1 effector conversion. Taken together, IL-23 and IL-1β signals are sufficient to induce Vγ6^+^ Tγδ17 cell plasticity by regulating the function of signal-dependent TFs including AP-1, STAT4, and NF-κB.

During homeostasis, many lymphocyte lineages with tissue-resident phenotypes express co-inhibitory receptors^50,86^. Here, we identify TIM-3 as a selective marker of type 1 effector-converted Vγ6^+^ ex-Tγδ17 cells with enhanced IFNγ production and proliferation following *S. typhimurium* infection. While PD-1 is expressed broadly by tissue-resident Tγδ17 cells in colon, lung, and uterus during homeostasis^7,43,51^, induction of TIM-3 is induced more selectively and is exclusive to Vγ6^+^ ex-Tγδ17 cells. Interestingly, TIM-3 is upregulated by Vγ4^+^ Tγδ17 cells in the lung tumor environment where it counteracts proliferation^51^. Co-inhibitory receptors are generally associated with inhibition of proliferation and effector functions, with TIM-3 often linked with exhausted CD8^+^ T cells^87^. However, consistent with our findings for Vγ6^+^ ex-Tγδ17 cells, both mouse skin CD8^+^ tissue-resident memory T (TRM) cells and human lung CD8^+^ TRM cells co-express PD-1 and TIM-3, yet proliferate in response to viral challenge^55^ and display superior functionality and proliferation^88^, respectively. Thus, expression of multiple co-inhibitory receptors is not always indicative of an exhaustion state and may instead reflect heightened functionality, especially for tissue-resident lymphocytes. The identification of cell surface markers for effector converted Vγ6^+^ ex-Tγδ17 cells such as TIM-3 will facilitate the evaluation of Tγδ17 cell plasticity in diverse contexts and genetic models.

Despite sharing a high degree of transcriptional similarity at homeostasis^43,51,89^, Vγ6^+^ and Vγ4^+^ Tγδ17 cells display divergent responses and capacities for plasticity following oral *S. typhimurium* infection. After infection, colonic Vγ6^+^ Tγδ17 cells undergo conversion to IFNγ single producers, robustly proliferate based on Ki-67 status, and upregulate TIM-3, whereas Vγ4^+^ Tγδ17 cells remain stable producers of IL-17A, retain a Ki-67^lo^ TIM-3^lo^ phenotype, and show negligible plasticity. Moreover, the downregulation of AP-1 TF expression post-infection— particularly of JunB and Fosl2 that have regulatory roles in restraining plasticity—is restricted to the Vγ6^+^ subset. Notably, divergent functionalities have also been reported in the lung, where opposing expression patterns for PD-1 and TIM-3 by Vγ6^+^ and Vγ4^+^ Tγδ17 cells result in their differential control by these co-inhibitory signals at steady state and in cancer^51^. Although, innately programmed, selective responses by Vγ6^+^ Tγδ17 cells to *S. typhimurium* infection may be driven by their unique TCR. In support of this view, the ligand for the semi-invariant Vγ6Vδ1 TCR is upregulated by bacterial infection and other inflammatory stimuli^56^ and the proliferative response of Vγ6^+^ Tγδ17 cells following *Listeria monocytogenes* challenge has been shown to be TCR-dependent^90^. Thus, Tγδ17 cell subsets have distinct regulatory wiring, likely due to their unique ontogenies^11^, TCRs, or cell surface receptors^43,51^, that result in differential responses and capacities for plasticity in the same environmental context. The conditions, if any, that elicit effector conversion of Vγ4^+^ Tγδ17 cells remain to be defined.

Taking our findings together, we have identified a core AP-1 regulatory axis that restricts type 1 plasticity in innate Tγδ17 cells. With the increasing appreciation of the role of Tγδ17 cells in health and disease, including the promotion of cancer through IL-17A production^32^, a better understanding of the mechanisms governing the flexibility of Tγδ17 cells could lead to potential therapies for diseases that benefit from enhanced type 1 and decreased type 3 effector functions.

## Methods

### Mice

Mice were housed under specific pathogen-free conditions and used in accordance with the Duke University Institutional Animal Care guidelines. Mice with floxed alleles for *Junb* (*Junb*^fl/fl^)^91^ and *Fosl2* (*Fosl2*^fl/fl^)^92^ were acquired from Erwin Wagner (CNIO, Spain); *Maf*^fl/fl^ mice were provided by C. Birchmeier (Max-Delbrück Center for Molecular Medicine, Germany)^93^, and *Bach2*^fl/fl^ mice were obtained from Tomohiro Kurosaki (Osaka University, Japan)^94^. RORgt-E2-Crimson reporter mice were kindly provided by Jinfang Zhu (NIH, US)^41^. The following commercially available strains were bred in our facility: *Rorc-*Cre mice (Stock 022791, Jackson), *Il17a*^Cre^ mice (Stock 016879, Jackson), *Rosa26^ZsGreen^*(Stock 007906, Jackson, referred to as *R26^ZSG^*), *Rosa26^tdtomato^* (Stock 007914, Jackson, referred to as *R26^TOM^*), and IFNg-IRES-eYFP (Stock 017581, Jackson). Adult mice were used between the ages of 8-16 weeks.

### Cell isolation from tissues

Lamina propria lymphocytes from the small and large intestine were isolated as previously described^80^. The same digestion protocol was applied to isolate immune cells from lung tissue. For female reproductive tract (FRT) processing, the vagina, uterine horn, and cervix were treated in the same manner as intestines^80^ except that the digestion time was extended to 60 minutes. Single cell suspensions of lymph nodes were made through mechanical disruption through a 40 μm cell strainer.

### Cell stimulation and Flow cytometry

To evaluate cytokine expression, *ex vivo* lamina propria lymphocytes or Tγδ17 cell cultures were stimulated for 4 h at 37°C in complete IMDM (IMDM supplemented with 10% FBS, 10 U ml^−1^ penicillin, 10 μg ml^−1^ streptomycin, 2 mM glutamine, 50 μg ml^−1^ gentamycin, and 55 μM β-mercaptoethanol) with phorbol 12-myristate 13-acetate (100 ng ml^−1^; Sigma-Aldrich) and ionomycin (375 ng ml^−1^; Sigma-Aldrich) in the presence of GolgiStop (BD) for the last 3 h. Additionally, to evaluate cytokine expression following IL-23 and IL-1β stimulation, *ex vivo* lamina propria lymphocytes were cultured for 20 h at 37°C in complete RPMI with IL-23 (25 ng ml^−1^ R&D Systems) and IL-1β (20 ng ml^−1^ Peprotech) in the presence of GolgiStop for the last 3h.

For all samples, Fc receptor blocking, surface stain, fixation/permeabilization, and TF staining was performed as described^80^. When sorting, cells were stained in PBS for 30 min at 4°C, washed, and resuspended in Ca^2+^/Mg^2+^-free PBS buffer containing 2mM EDTA and 0.5% BSA. Sorting was performed using a MoFlo Astrios or MoFlo XDP cell sorter (Beckman Coulter). In all experiments, a fixable viability dye (eBiosciences) was used to exclude dead cells. Antibodies were purchased from Biolegend (CD127, Vγ4, PD-1), BD Biosciences (CCR6, CD3), and eBiosciences (CD45, RORγt, CD3, CD11b, Ter1119, B220, CD19, T-bet, CD90.2, CD90.1, IL-17A, IFNγ, TCRβ, γδTCR, TIM-3, NKp46, T-bet). The Vγ6 antibody was kindly provided by Dr. Yasunobu Yoshikai (Kyushu University, Japan)^95^.

### Expansion of T**γδ**17 cells from mesenteric LN

The protocol for selective expansion of Tγδ17 cells from mLN was adapted from Mckenzie et al^37^. Complete RPMI supplemented with 1X NEAA and 1X Glutamax was used for cultures. 96-well round bottom plates were coated with 1 μg ml−1 of γδTCR antibody (clone GL3). Plates were incubated at 37°C for 4 hours, and thereafter, washed twice with PBS. mLN cells were cultured at 2 x 10^5^ cells/well in the presence of IL-23 (4 ng ml−1), IL-1β (5 ng ml−1), and anti-IFNγ (10 μg ml−1). 3 days later, cells were harvested, washed, and re-plated at 100,000 cells/well in a flat bottom 96-well plate in the presence of IL-23 (4 ng ml−1), IL-1β (5 ng ml−1), and anti-IFNγ (10 μg ml−1). On day 5, cells were transduced with retroviral supernatants containing hexadimethrine bromide (6.66 ng ml−1; Sigma-Aldrich) for 120 minutes at 2400rpm and 30°C. On day 6, media was exchanged for new media with IL-7 (20 ng ml−1) and anti-IFNγ (10 μg ml−1). On day 9, cells were harvested for flow cytometry or were stimulated with PMA/Ionomycin in the presence of Golgistop for 4h for analysis of cytokine production.

### Retroviral Gene Transfer

Retroviral constructs were generated by cloning complementary DNA for the TF of interest into murine stem cell virus (MSCV)-Thy1.1 5′ of the internal ribosomal entry site, allowing bicistronic expression with cell surface Thy1.1. Dominant negative (DN) Fosl2 overexpression vector was generated by truncating the carboxyl-terminus (amino acids 207-326)^68^. Retroviral supernatants were generated by transfection of retroviral constructs into the Plat-E producer cell line^96^ using Lipofectamine 2000 reagent (ThermoFisher Scientific) and collection of culture media after 48 h. Retroviral supernatants were filtered with 0.45-μm filters before use.

### Intestinal *S. typhimurium* infection

Mice in both naïve and infection groups were fasted for four hours, followed by oral gavage with 20 mg of Streptomycin (Sigma-Aldrich) in 200ul of PBS. 20 hours later mice were fasted again for four hours. For infection, 5×10^7^ colony forming units (CFU) of *Salmonella typhimurium* (Strain SL1344) in 200ul PBS were orally gavaged. 200ul of PBS was gavaged for naïve controls. Mice were monitored for weight loss. Approximately 96 hours post infection, mice were euthanized and tissues were harvested. SL1344 was kindly provided by Dr. Soman Abraham.

### Intestinal *C. rodentium* infection

Mice were fasted for four hours, followed by oral gavage with 2×10^9^ (CFU) of *Citrobacter rodentium* Schauer et al. (ATCC 51459^97^, DBS100) in 200ul of PBS to induce colitis. 200ul of PBS was gavaged for naïve controls. All mice were monitored for weight loss over the course of infection. Mice were euthanized 13 days post infection and coLP tissue was harvested.

### Cell sorting for 10X Genomics single cell multiome assay

Total γδ T cells (CD90^+^ CD3^+^ γδTCR^+^) were isolated and sorted from the coLP of 96 hour *S. typhimurium*-infected mice (sample pooled from 5 mice), and “naïve” Streptomycin-only treated mice (sample pooled from 4 mice). Total coLP ILCs (CD3^−^ CD127^+^ CD90^+^) were also sorted from the same infected and naïve samples, and γδ T cells and ILCs were combined for multiome assay. The ILC data are not included in this manuscript and will be reported elsewhere.

### Single cell 10X Genomics multiome library preparation

Nuclei were isolated from cells using a detergent based lysis buffer to interrupt the cell membrane and release the contents. After nuclei were purified through centrifugation and washes, they were treated with transposase and partitioned into nanoliter-scale Gel Beads-in-emulsion (GEMs) using the Chromium Controller (10x Genomics, Pleasanton, CA, USA). The transposase enters the nuclei, preferentially fragments DNA in regions of open chromatin, and simultaneously adds adapter sequences to the ends of the DNA fragments. The transposed nuclei were then loaded onto a microfluidic chip along with ATAC + GEX Gel Beads, Master Mix, and partitioning oil to generate oil emulsions. The GEMs were then transferred from the chip onto a thermocycler for incubation. During this incubation, Gel Beads dissolve, releasing oligonucleotides containing an Illumina TruSeq Read 1 (read 1 sequencing primer), 16 nt 10x Barcode, 12 nt unique molecular identifier (UMI), and a 30 nt poly(dT) sequence that enables production of barcoded, full-length cDNA from poly-adenylated mRNA for the gene expression (GEX) library. Oligos containing a Spacer sequence, Illumina® P5 sequence, a 16 nt 10x Barcode that enables barcode ligation to transposed DNA fragments for the ATAC library were also released. Both oligos were mixed with the nuclei lysate containing transposed DNA fragments, mRNA, and reagents for reverse transcription (RT). Incubation of the GEMs produced 10x Barcoded DNA from the transposed DNA (for ATAC) and 10x Barcoded, full-length cDNA from poly-adenylated mRNA (for GEX). After incubation, the GEMs were broken and pooled fractions were recovered and amplified via PCR to fill gaps and to generate sufficient mass for library construction. The pre-amplified product was used as input for both ATAC library construction and cDNA amplification for gene expression library construction. The P7 Illumina adapter sequence and a unique sample index were added during ATAC library construction via PCR. The pre-amplified, Barcoded, full-length cDNA product was also amplified once again via PCR to generate sufficient mass for gene expression library construction. This amplified product was then used for a standard NGS library prep. Briefly, enzymatic fragmentation and size selection were used to optimize the cDNA amplicon size. P5, P7, i7 and i5 sample indexes, and TruSeq Read 2 (read 2 primer sequence) were added via End Repair, A-tailing, Adaptor Ligation, and index PCR. The final GEX and ATAC libraries were then sequenced on an Illumina Novaseq SP (Illumina, San Diego, CA, USA). Single cell Multiome ATAC + Gene Expression v1 chemistry was used. The naïve sample had 6,399 cells sequenced and the *S. typhimurium* sample had 6,510 cells sequenced.

### Single cell 10X Genomics multiome data analysis

The 10X Genomics Cell Ranger pipeline (cellranger-arc-2.0.0) was used to generate FASTQ files for the ATAC^98^ and Gene expression^99^ libraries. FASTQ files were aligned to the mouse mm10 genome. The ZsGreen (*Zfp506*) sequence was added to the reference mouse genome ‘mm10 Reference - 2020-A-2.0.0’ before calling cellranger-arc-2.0.0. Downstream analysis was performed in RStudio using Seurat^42^ Version 4.1. Naïve and *S. typhimurium* objects were merged into one object after processing. 11,547 cells passed QC. The SCTransform^100^ function in Seurat^42^ was used for normalization and regression of cell cycle scoring. Principal component analysis (PCA) was performed on all genes, and the elbow method was used to determine the number of principal components to be utilized. Thirty principal components were used for clustering and dimensionality reduction using FindNeighbors and RunUMAP in Seurat, which identified 22 distinct cell clusters. The RNA assay in the Seurat object was then multiplied by 10,000 and log-transformed before cluster-specific genes were identified. Clusters were manually labeled based on top genes expressed using the FindConservedMarkers function. Fifteen clusters and 9,950 cells remained after contaminating myeloid, B cells, and epithelial cells were removed from the object. Re-clustering of only γδ T cells led to 2,881 cells dividing into 14 clusters. FindMarkers was used for differential expression analysis using the MAST statistical test. Gene expression, gene signature scores, and clustering results were all visualized by embedding cells in dimensionally-reduced UMAP space. Gene signature scores were calculated using the AddModuleScore function in Seurat and the Nebulosa^101^ package was used to produce density plots in UMAP space. Motif analysis was done using the Signac^102^ package. The per-cell motif activity score was calculated using chromVAR^103^ to identify differentially active motifs between clusters associated with variation in chromatin accessibility.

### RNA velocity analysis

The spliced and unspliced mRNA count matrix was generated utilizing Velocyto^46^, which processed the raw BAM and loom data from single cell RNA sequencing (scRNA-seq). Information about cell cluster number, PCA coordinates and WNN UMAP coordinates from the Seurat object was transferred to python-based AnnData format for further analysis with Scanpy^104^. UniTVelo, a method employed for modeling dynamics through flexible transcription activities, was utilized to infer RNA velocity^47^. Parameters for UniTVelo configuration were set to linear regression R-squared on extreme quantile, none root cell cluster specified, and using a unified-time mode. To model the rate of change of spliced and unspliced mRNA counts, 2000 highly variable genes were selected for each gene in every cell, facilitating the inference of directionality and speed of gene expression changes over time. Subsequently, RNA velocity vectors were estimated for individual cells. RNA velocity was further visualized using scvelo.pl.velocity_embedding_stream to predict future cell states and trajectories within the dataset, employing WNN UMAP for visualization.

### Simulating molecular dynamics in pseudo-time trajectory of V**γ**6 plasticity

The TSCAN and Monocle3 pseudo-time analyses were utilized to delineate the cell trajectory, reflecting the anticipated linear or branching biological topologies^48,49,105^. The roots of the trajectories were manually selected from the naïve Vγ6 cluster (cluster 0). Subsequently, state transition inference prediction (STIP) was leveraged to discern gene expression patterns and transcription factor (TF) motif activities along this trajectory^106^. STIP identifies genes with significant expression changes along the pseudo-time trajectory, averages their expression across all cells, and standardizes it to a mean of zero and a variation of one. Then, it estimates the pseudo-time point at which the standardized expression is zero for each gene (zero point), retaining only those with one or two zero points. Genes are reordered based on their expression patterns and the location of the zero point within each pattern. The same process applies to the TF motif activities computed by chromVar using the JASPAR curated motif database^103,107^. Nonparametric regression curves for mRNA expression and TF motif activity along the pseudo-timeline were computed by fitting the normalized matrix to a generalized additive model (GAM)^48^. Regression curves were further plotted for visualizing changes throughout the plasticity trajectory.

### Gene Regulatory Network (GRN Regulon) Analysis

The SCENIC (Single-Cell rEgulatory Network Inference and Clustering) computational pipeline was deployed to infer gene regulatory networks within single cell sub-clusters (cluster 0, 6, 7, 9), facilitating the identification of pivotal regulatory drivers governing cellular states^57,108^. The SCENIC pipeline was executed in R using default parameters, leveraging the normalized single cell mRNA gene expression matrix generated by Seurat as input data. The Scenic object was initialized to settings using mouse-specific databases for RcisTarget. RcisTarget is a function that scores the motifs in the promoter of the genes (up to 500bp upstream the TSS), and in the 20kb around the TSS (+/-10kbp). Following that, GENIE3^109^ conducted co-expression analysis to discern gene regulatory elements and predict their target genes, capitalizing on the co-expression patterns of transcription factors (TFs) and their putative targets across individual cells. Regulon activities were visually depicted and plotted for comprehensive analysis.

### Statistical Analysis

Apart from single cell RNA/ATAC-seq, all datasets were analyzed using either GraphPad Prism 9 or 10. Flow cytometry phenotyping data underwent analysis employing two-tailed unpaired and paired Student’s t-tests, Ordinary One-Way ANOVA, and Ordinary Two-Way ANOVA. Samples containing fewer than 50 cells in the final gate for flow cytometric analysis were omitted from further analyses. Grubbs’ tests were utilized to detect and remove individual outliers in summary data.

## Acknowledgements

We thank Eunchong Park for assistance with gut preps. We also thank the Molecular Genomics Core for processing and running the 10X Genomics Multiome samples. We acknowledge the expert assistance of Lynn Martinek with flow cytometry sorting, and Scott Langdon at the Duke DNA Sequencing Facility. Lastly, we thank Dr. Yasunobu Yoshikai (Kyushu University, Japan) for sharing the Vγ6 antibody. This work was funded by NIH grant R01 GM115474 and P01 AI102853. M.E.P was supported by F31 AI152457. N.U.M was supported by F31 AI181082.

---

**Extended Data Fig. 1.**
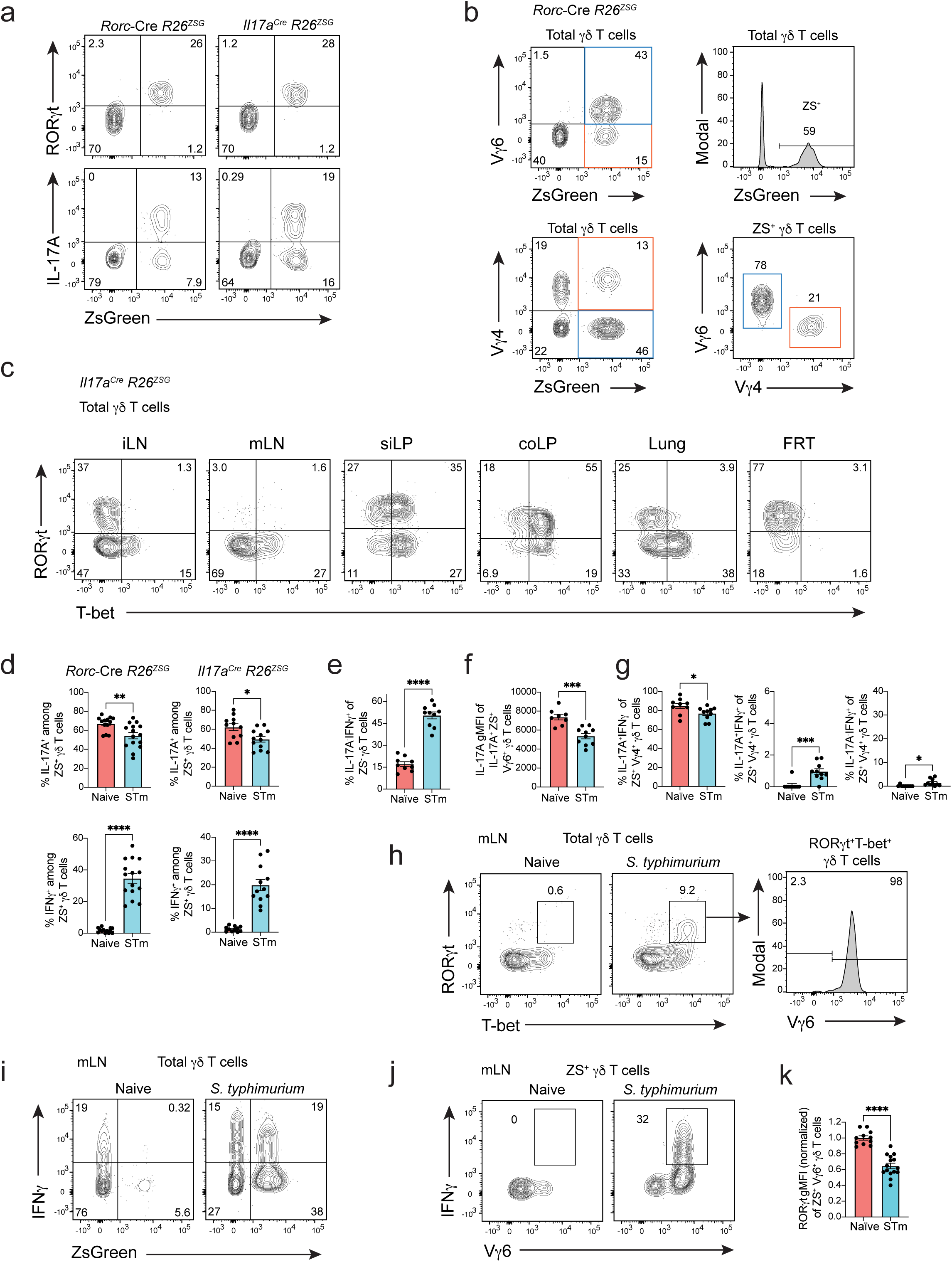
Tγδ17 cells are stable at steady state and Vγ6^+^ Tγδ17 cells are plastic after *S. typhimurium* in mLN and coLP. (**a**) Flow cytometric analysis of coLP of *Rorc*-Cre *R26^ZSG^*and *Il17a*^Cre^ *R26^ZSG^* fate-mapping mice. RORγt, IL-17A, and ZS-green expression gated on total γδ T cells. Flow plots representative of more than three independent experiments, n >10. (**b**) Flow cytometric analysis of coLP of *Rorc*-Cre *R26^ZSG^* measuring Vγ6, Vγ4, and ZS-green expression gated on total total γδ T cells (left and top right). Bottom right is gated on ZS^+^ γδ T cells. Flow plots representative of more than three independent experiments, n > 9. (**c**) Flow cytometric analysis of total γδ T cells from coLP of *Il17a*^Cre^ *R26^ZSG^* fate-mapping mice for RORγt and T-bet expression. Flow plots representative of more than three independent experiments (n =8) except lung is n=5 from one experiment. (**d**) Summary data of percentage of IL-17A and IFNγ in naïve and *S. typhimurium* (STm) coLP for *Rorc*-Cre *R26^ZSG^*and *Il17a*^Cre^ *R26^ZSG^* fate-mapping mice. Gated on ZS^+^ γδ T cells. Each summary graph pooled from three independent experiments n=11 or more. (**e**) Summary data of percentage of IL-17A^−^IFNγ^+^ in naïve and *S. typhimurium* (STm) coLP for *Il17a*^Cre^ *R26^ZSG^* fate-mapping mice gated on ZS^−^ γδ T cells. Each summary graph pooled from two independent experiments, n=9 or more. (**f**) Summary data of IL-17A gMFI on IL-17A^+^ Vγ6^+^ ZS^+^ γδ T cells. (**g**) Summary data of percentage of IL-17A^+^IFNγ^−^, IL-17A^+^IFNγ^+^, IL-17A^−^IFNγ^+^ in naïve and *S. typhimurium* (STm) coLP for *Il17a*^Cre^ *R26^ZSG^* fate-mapping mice gated on Vγ4^+^ ZS^+^ γδ T cells. Each summary graph pooled from two independent experiments, n=9 or more. (**h**) Flow cytometric analysis of mLN of *Rorc*-Cre *R26^ZSG^* mice gated on total γδ T cells (left) for RORγt and T-bet expression and Vγ6 expression in RORγt^+^T-bet^+^ γδ T cells (right). Representative flow plot from three independent experiments, n > 10 mice. (**i**) Flow cytometric analysis of IFNγ production by ZS^+^ γδ T cells in mLN in naïve and *S. typhimurium* infected *Rorc*-Cre *R26^ZSG^* mice. Representative flow plot from three independent experiments, n > 10 mice. (**j**) Flow cytometric analysis of Vγ6 versus IFNγ production by ZS^+^ γδ T cells in mLN in naïve and *S. typhimurium* infected *Rorc*-Cre *R26^ZSG^*mice. Representative flow plot from three independent experiments, n > 10 mice. (**k**) Summary data pooled from three independent experiments for RORγt expression among Vγ6^+^ ZS^+^ γδ T cells for ZS^−^ and ZS^+^ populations from naïve and (STm) *S. typhimurium* infected *Rorc*-Cre *R26^ZSG^* mice, n > 10 mice. All results represent mean ± s.e.m. *P < 0.05; **P < 0.01; ****P < 0.0001; ns, not significant (two-tailed unpaired Student’s t-test). Numbers in flow plots represent percentages of cells in the gate.

**Extended Data Fig. 2.**
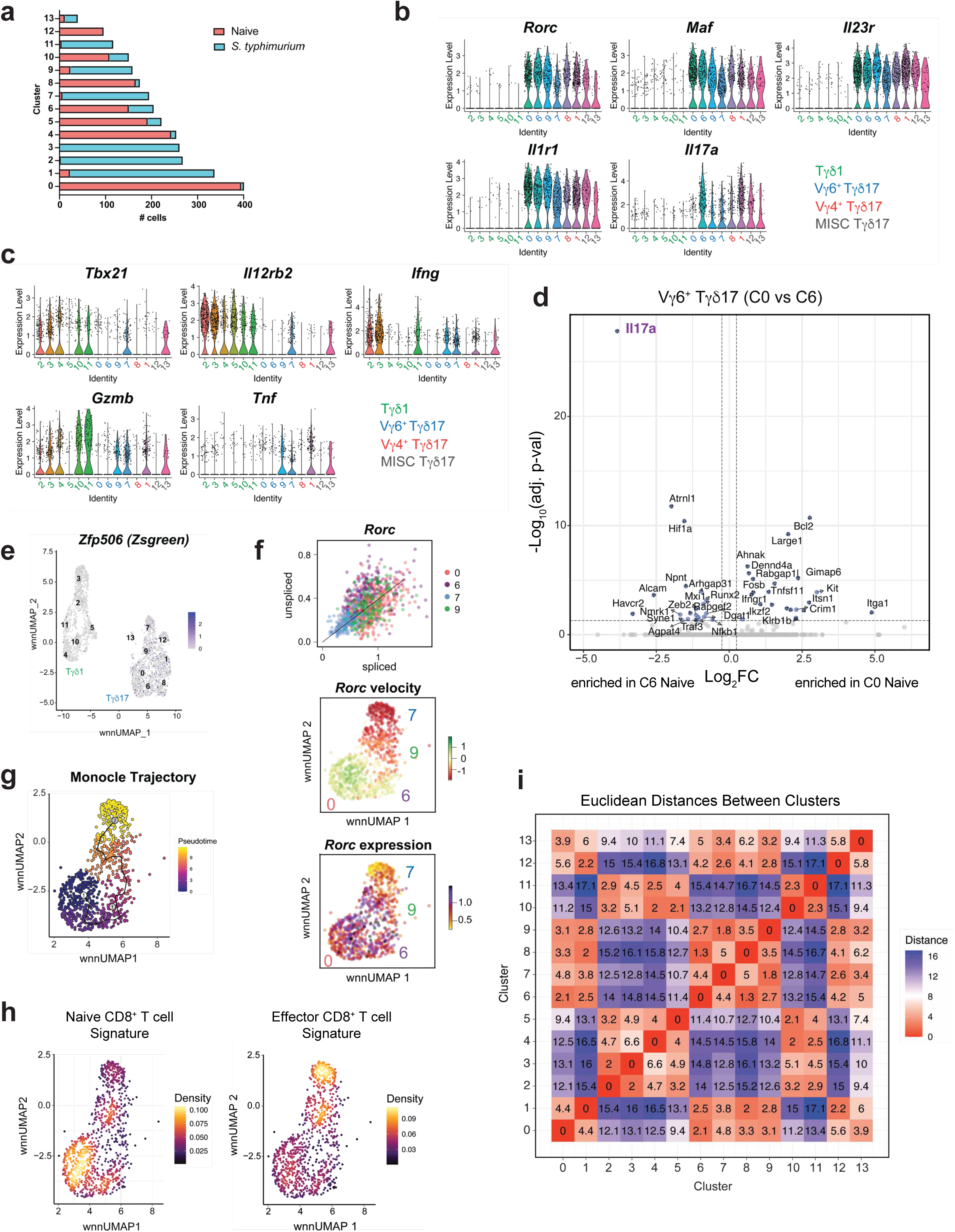
Single cell multiome characterization of γδ T cells and Vγ6^+^ Tγδ17 cell trajectories. (**a**) Barplot of number of cells in each cluster and from which condition for total γδ T cells. (**b,c**) Violin Plots of type 3 and 1 genes for γδ T cells clusters. (**d**) Volcano plot for cells from naïve condition for C0 vs C6 Vγ6^+^ Tγδ17 cells with blue dots indicating significance based on p-val adj < 0.05 and log_2_FC > 0.25; FC, fold change. (**e**) *zFP506* (ZsGreen) FeaturePlot for γδ T cell clusters. (**f**) Plot of per cell unspliced versus spliced *Rorc* transcript, RNA velocity for *Rorc*, and *Rorc* expression FeaturePlot all for Vγ6^+^ Tγδ17 clusters C0, C6, C9 and C7. (**g**) Monocle 3 trajectory of Vγ6^+^ Tγδ17 clusters C0, C6, C9 and C7. (**h**) CD8^+^ T cell gene signatures (GSE9650) projected onto UMAP space with Seurat’s AddModuleScore function. Nebulosa Plot Density plot displaying enrichment of gene signature. (Left) Genes upregulated in naive vs effector CD8^+^ T cells. (Right) Genes downregulated in naive vs effector CD8^+^ T cells. (**i**) Euclidian distances between γδ T cell clusters based on WNN UMAP.

**Extended Data Fig. 3.**
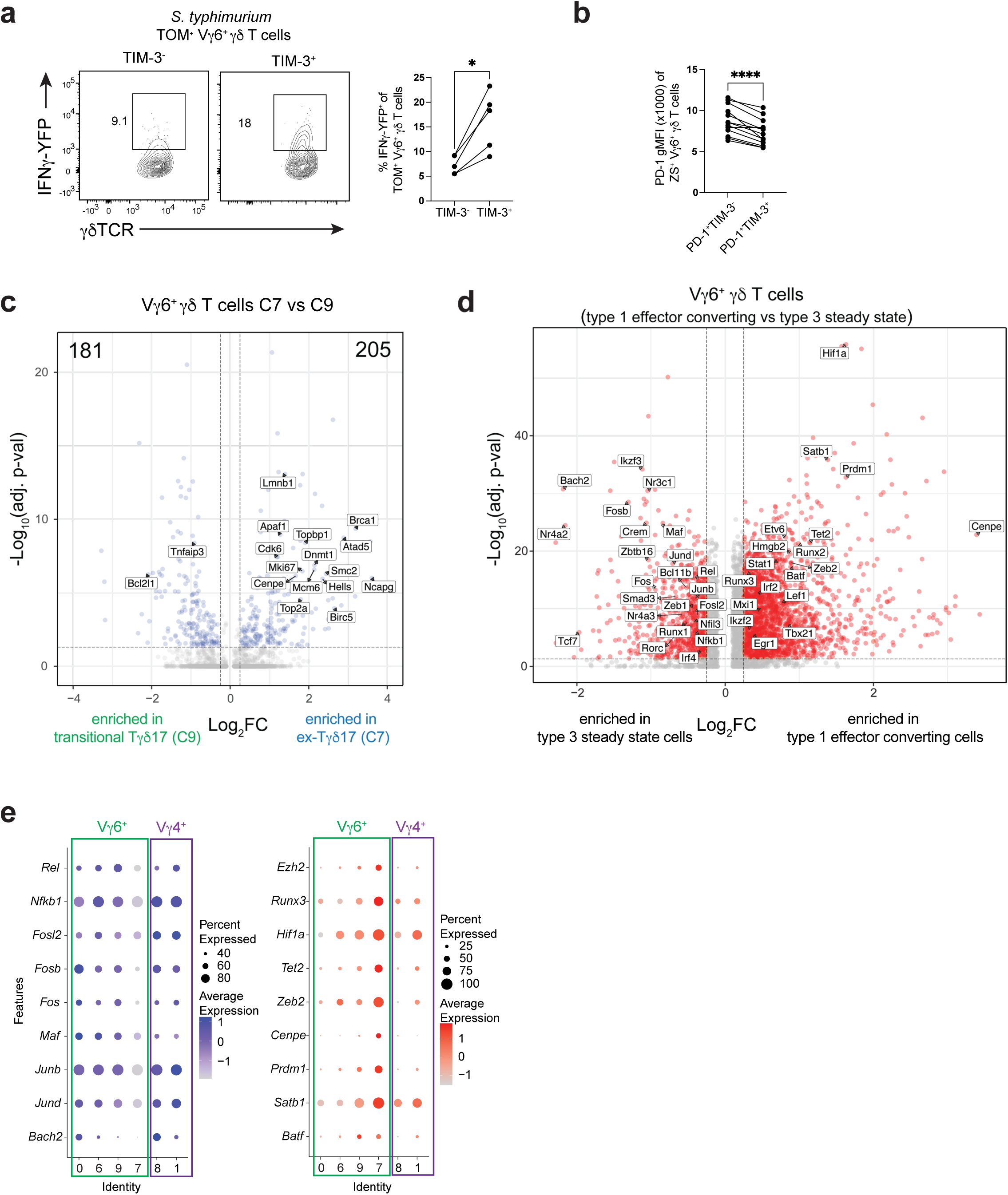
Effector converted Vγ6^+^ Tγδ17 cells have distinct transcriptional profiles compared to steady state. (**a**) IFNγ-YFP expression in fate-mapped TIM-3^−^ or TIM-3^+^ Vγ6^+^ TOM^+^ γδ T cells from naïve and *S. typhimurium* infected *Il17a*^Cre^ *R26^TOM^* IFNγ-YFP mice. Summary plot from one experiment n=5 mice. (**b**) Summary data for PD-1 gMFI in PD-1^+^TIM-3^−^ or PD-1^+^TIM-3^+^ Vγ6^+^ ZS^+^ γδ T cells compiled from three experiments, n=13 mice. (**c**) Volcano plot for differentially expressed genes between C7 vs C9 Vγ6^+^ Tγδ17 cells with blue dots having p-val adj < 0.05 and log2FC > 0.25. (**d**) Volcano plot of differentially expressed transcription factors in type 1 converting Vγ6^+^ γδ T cell clusters (C7+C9) compared to type 3 steady state clusters (C0+C6) with red dots having p-val adj < 0.05 and log_2_FC > 0.25; FC, fold change. (**e**) DotPlot for Tγδ17 cell clusters for select transcriptional regulators downregulated (left) or upregulated (right) in Vγ6^+^ Tγδ17 cells with Vγ4^+^ Tγδ17 cells for comparison. All results represent mean ± s.e.m. Paired t-test for (**a-b**). *P < 0.05; ****P < 0.0001; ns, not significant (two-tailed paired Student’s t-test). Numbers in flow plots represent percentages of cells in the gate.

**Extended Data Fig. 4.**
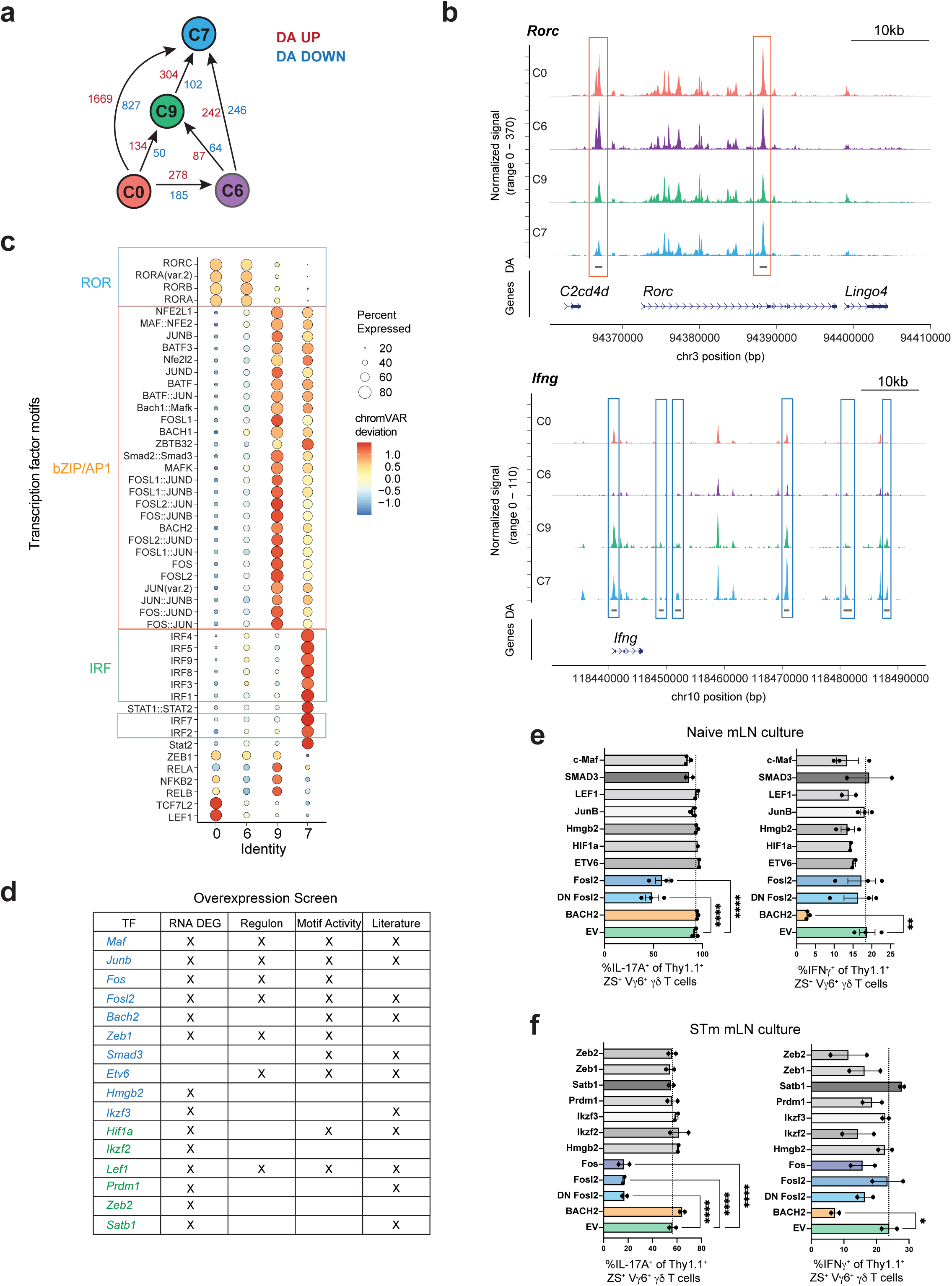
BACH2 and AP-1 TFs regulate Vγ6^+^ Tγδ17 plasticity in vitro. (**a**) Schematic of Vγ6^+^ Tγδ17 cluster 0, 6, 9, and 7 for number of regions differentially accessible (DA) (p < 0.05) with regions increasing (UP, red) and decreasing (DOWN, blue) in accessibility. (**b**) Pseudobulk scATAC-seq CoveragePlots for *Rorc* and *Ifng* loci for Vγ6^+^ Tγδ17 cluster 0, 6, 9, and 7. Rectangle highlights regions with significant differential accessibility (p < 0.05) shown for decreasing (orange) or increasing (blue) in accessibility. (**c**) Motif activity dot plot of Vγ6^+^ Tγδ17 clusters using chromVAR with colored boxes highlighting specific TF families. (**d**) TFs in Vγ6^+^ Tγδ17 cell overexpression screen. X’s in RNA DEG column means the TF of interest is DEG at some point along trajectory. X in Regulon column means the TF shows up in regulon analysis. X in Motif Activity column means TF has differential motif activity (chromVAR) during conversion. X in Literature column means TF is implicated in type 3 lymphocyte regulation. Blue TFs predicted to stabilize type 3 program and green TFs predicted to promote type 1 conversion. (**e**) Flow cytometric analysis of cytokine production from 9 day Tγδ17 mLN culture. Gated on transduced Vγ6^+^ (Vγ4^−^) Thy1.1^+^ ZS^+^ γδ T cells from steady state *Il17a^Cre^R26^ZSG^*mice after 4 hours PMA/Ionomycin stimulation. Summary graph pooled from two independent experiments. (**f**) Same as in e but from *S. typhimurium* infected *Il17a^Cre^R26^ZSG^* mice. Summary graph from one independent experiment. Statistical analyses included Ordinary one-way ANOVA tests for (**e,f**). Results represent mean ± s.e.m. \**P* < 0.05; \*\**P* < 0.01; ***P < 0.001; \*\*\*\**P* < 0.0001; DEG, differentially expressed gene; ns, not significant.

**Extended Data Fig. 5.**
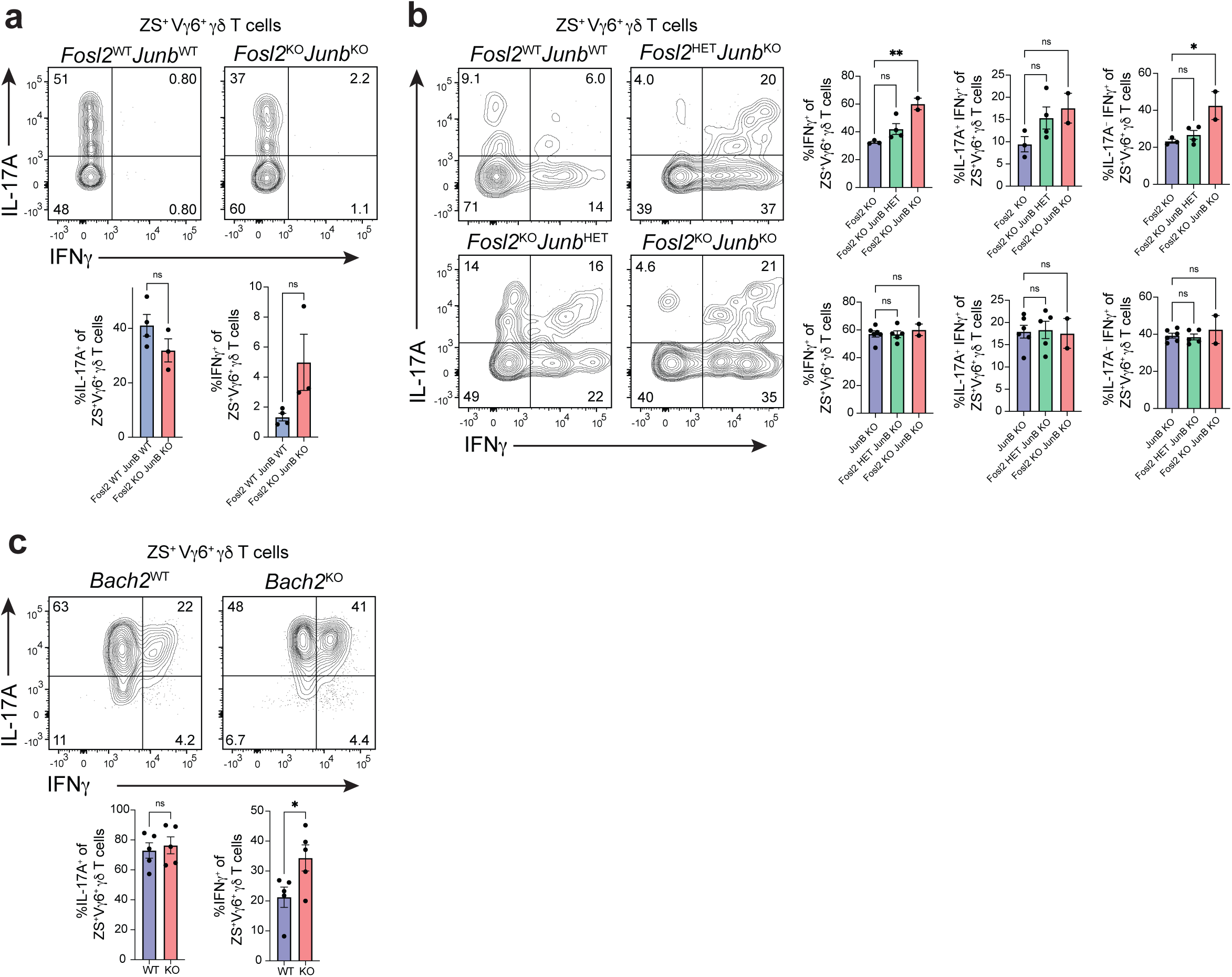
JunB plays a more prominent role than Fosl2 in Vγ6^+^ Tγδ17 cell plasticity. (**a-b**) Flow cytometric analysis was performed on colonic ZsGreen^+^ Vγ6^+^ Tγδ17 cells from mice with compound *Junb* and *Fosl2* conditional deletions on the *Il17a^Cre^R26^ZSG^* deleter background at steady state (*TF^+/+^, TF*^WT^; *TF^fl/+^, TF*^HET^; *TF^fl/fl^, TF^KO^*): (**a**) Representative flow cytometric analysis of the frequency of IL-17A and IFNγ producing cells following 4 h PMA/ionomycin stimulation. (*n* = 3-4 mice/genotype; two independent experiments). (**b**) Representative flow cytometric analysis and summary plots of the frequency of IL-17A and IFNγ producing colonic ZS^+^ Vγ6^+^ Tγδ17 cells at steady state following 20-h stimulation with IL-23 and IL-1β. (*n* = 2-11 mice/genotype; two independent experiments). (**c**) Flow cytometric analysis was performed on ZS^+^ Vγ6^+^ from mLN of naïve *Bach2^+/+^Il17a^Cre^R26^ZSG^*(*Bach2*^WT^) and *Bach2^fl/fl^Il17a^Cre^R26^ZSG^* (*Bach2*^KO^) mice on day 9 of Tγδ17 mLN culture. Summary plots of the frequency of IL-17A and IFNγ producing cells following 4 h PMA/ionomycin stimulation (*n* = 5 mice/genotype; three independent experiments). (**a-c**) Gating was performed on fate-mapped Vγ4^−^ Tγδ17 cells (CD3ε^+^γδTCR^+^TCRβ^−^ZS^+^Vγ4^−^). Statistical analyses include Two-tailed unpaired Student’s t-tests for (**a,c**) and an Ordinary one-way ANOVA test for (**b**). Results represent mean ± s.e.m. \**P* < 0.05; \*\**P* < 0.01; ns, not significant. Numbers in flow plots represent percentages of cells in the gate.

